# Genetic determinants of *Brevundimonas*-mediated attenuation of plant pattern-triggered immunity

**DOI:** 10.64898/2026.10.27.762230

**Authors:** Chao Liao, Dinie Zheng, Anthea Zhang, Jay C. Georges, Jonathan M. Conway

## Abstract

Root-associated commensal bacteria from diverse taxonomic groups can attenuate plant pattern-triggered immunity (PTI), yet the bacterial mechanisms underlying immune attenuation remain poorly understood. *Brevundimonas* sp. MF374 is a root-associated Alphaproteobacterium previously identified as a strong attenuator of flg22-triggered immunity. Here, to identify the genetic determinants underlying this phenotype, we conducted transposon mutagenesis and screened 4,032 MF374 mutants for defects in immune attenuation. Deletion and complementation of candidate loci identified *tonB1* (BLR19_RS15045) as the strongest determinant of immune attenuation, with *ppiD* (BLR19_RS02880) making a weaker contribution. A panel of GUS reporter assays designed to distinguish among different mechanisms of attenuation showed that intact MF374 cells directly reduced the immune-eliciting activity of flg22 in a *tonB1*-dependent manner. TonB1 also contributed to attenuation of responses triggered by a distinct flg22 sequence and the endogenous plant elicitor AtPEP1. Together, these findings implicate TonB-dependent transport in attenuation of flg22-induced immunity by MF374 and reveal a previously uncharacterized mechanism of immune attenuation by a root-associated commensal.

## Introduction

Plant-microbe interactions play a critical role in determining plant fitness and shaping microbial community assembly. The plant innate immune system enables defense against microbial invasion by detecting conserved Microbe-Associated Molecular Patterns (MAMPs), including flagellin-derived flg22, elongation factor Tu-derived elf18, chitin, and lipopolysaccharides (*1–7*) among others. Related immune responses can also be triggered by endogenous Damage-Associated Molecular Patterns (DAMPs), including Plant Elicitor Peptide 1 (PEP1) (*8*, *9*). These immune elicitors are perceived by Pattern Recognition Receptors (PRRs) localized at the plant cell surface, initiating Pattern-Triggered Immunity (PTI) through receptor activation and downstream immune responses including reactive oxygen species (ROS) burst, apoplastic pH and ion flux changes, transcription of defense-associated genes including for secondary metabolite production (*10–12*). Activation of these immune responses is frequently accompanied by reduced plant growth, a relationship commonly described as the growth-defense trade-off (*13–16*).

Certain plant-associated commensal bacteria can attenuate plant immune activation (*13*, *17–21*). Such immune attenuation can influence plant growth and microbial community assembly by reducing immune-associated plant growth inhibition and facilitating colonization by commensal and beneficial microbes (*18*, *19*). Recent studies have identified that this activity is widespread among taxonomically diverse rhizosphere commensals, with approximately 40% of strains exhibiting immune attenuation activity in tested isolate collections (*13*, *17*, *18*). The diversity of strains possessing this phenotype suggests that multiple mechanisms may underlie immune attenuation, yet only a limited number of mechanisms have been characterized to date and span a range of distinct biological processes. *Pseudomonas capeferrum* WCS358 and *P. simiae* WCS417 can attenuate flg22-triggered immune responses by modifying environmental pH through gluconic acid production (*18*, *20*). A distinct mechanism was identified in *Dyella japonica* MF79, which secretes the subtilase IssA through the type II secretion system; IssA cleaves flg22 and other peptide immune elicitors to attenuate immune activation (*17*, *19*). IssA homologs are also present in other plant-associated commensals and are particularly conserved among members of the order Xanthomonadales (*17*). More recently, mutant screening in *Rhodanobacter* sp. R179 identified DssA and DssB, an ABC transporter permease and TonB-dependent transporter, respectively, as determinants of flg22-triggered immune attenuation in this strain, although the substrates and mechanistic roles of these transport systems remain unresolved (*21*). Cell-free supernatant from R179 also eliminates several immunogenic peptides through a separate extracellular peptidase activity that is retained in a Δ*dssAB* mutant (*21*). Together, these studies indicate that different rhizosphere commensals employ distinct strategies to attenuate plant immune activation, suggesting that additional uncharacterized mechanisms are likely to exist in other bacteria.

*Brevundimonas* sp. MF374 is a root-associated Alphaproteobacterium that is phylogenetically distinct from the Gammaproteobacterial commensals in which immune attenuation mechanisms have been characterized to date. Teixeira et al. (*19*) first identified MF374 as a potent attenuator of flg22-triggered immunity across root growth inhibition (RGI) and β-glucuronidase (GUS) reporter plant assays, and analyzed the plant transcriptome under MF374 and flg22 treatment. Our subsequent screening of a larger number of strains from the same isolate collection further demonstrated robust attenuation by MF374 (*17*). Despite this strong and reproducible attenuation of flg22-induced immunity, the molecular mechanism underlying this phenotype remains unknown.

In this study, we constructed transposon mutant libraries in MF374 and screened 4,032 mutants using an Arabidopsis GUS reporter assay, identifying five candidates defective in attenuation of flg22-triggered immunity. We subsequently validated the role of *tonB1* (one of four *tonBs* in MF374 genome annotated by NCBI RefSeq), *ppiD*, and *trpEXGDC* in this attenuation phenotype using genetic deletion and complementation. This analysis identified *tonB1* as the strongest determinant of MF374-mediated immune attenuation, with *ppiD* making a weaker contribution, while the phenotype associated with *trpEXGDC* was largely attributable to defects in bacterial growth and fitness. Using a series of modified GUS reporter assays to distinguish cell-associated, plant-mediated, and secreted mechanisms of attenuation, we found that interaction of flg22-*Pa* (derived from *Pseudomonas aeruginosa*) with intact MF374 cells substantially reduced its subsequent immune-eliciting activity, and that this phenotype was strongly dependent on *tonB1*. The contribution of *tonB1* also extended to attenuation of responses triggered by a distinct flg22 sequence, flg22-*Dja* (derived from *Dyella japonica* MF79), and the endogenous plant elicitor AtPEP1. Together, these results reveal a previously uncharacterized TonB-dependent mechanism in which direct interaction with MF374 cells removes immune-eliciting activity from the extracellular environment to attenuate plant immune activation.

## Results

### An RB-TnSeq screen identified MF374 mutants impaired in attenuation of flg22-triggered immune activation

To identify the MF374 genes involved in attenuation of immune activation triggered by flg22-*Pa*, we constructed RB-TnSeq mutant libraries in MF374 (**Figure 1A**) using mariner and Tn5 transposon vectors, pTGG31 (*22*) and pDZ003, respectively. Both libraries showed broad genome-wide insertion coverage with an average of 1.3 Tn insertions per 10 bp (**Figure S1**). In total 2,900 of 2,947 genes were identified to have Tn insertions, thus there were 47 genes that were not inserted with RB-Tn, indicating these genes are essential genes. Because the Tn5 library generated with pDZ003 contained more mapped unique barcodes than the mariner library generated with pTGG31 (**Figure S1D**, **Table S1**), the pDZ003 mutant library was selected for functional screening using the *pCYP71A12*::*GUS* reporter assay (**Figure 1A**).

**Figure 1.**
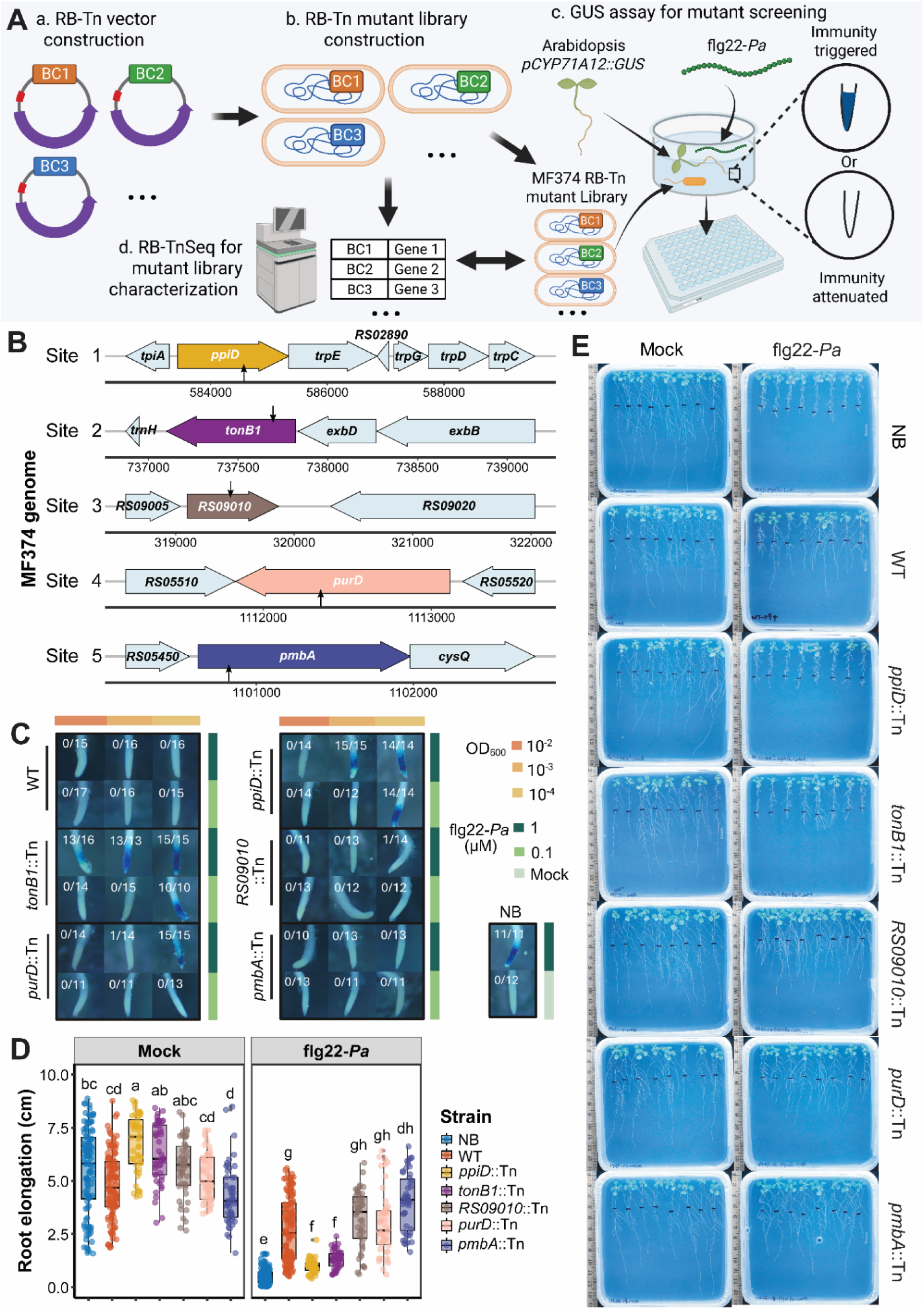
A transposon mutant screen identifies *tonB1* and *ppiD* as genetic determinants of immune attenuation by *Brevundimonas* sp. MF374. **(A)** Schematic of randomly barcoded transposon (RB-TnSeq) mutant library construction and screening for loss of immune attenuation using the *Arabidopsis thaliana pCYP71A12*::*GUS* line. BC: barcode. **(B)** Genomic locations (black arrows) of transposon insertions in five candidate mutants in the MF374 genome. **(C)** GUS reporter assays of MF374 wild type (WT) and candidate mutants at initial OD_600_ of 10^-2^ to 10^-4^ and flg22-*Pa* concentrations of 0.1 or 1 µM in *A. thaliana pCYP71A12*::*GUS*. Fractions indicate the number of blue-stained root tips relative to the total number examined across three biological replicates. **(D)** Root elongation of *A. thaliana pWER*::*FLS2-GFP* seedlings 9 days after inoculation with no bacteria (NB), wild type (WT), or mutant strains at OD_600_ = 0.01, with or without 1 µM flg22-*Pa.* Different letters across both mock and flg22-*Pa* panels indicate significant differences (*P* < 0.05; Kruskal-Wallis test followed by Dunn’s test with the Benjamini-Hochberg method for *P*-value adjustment of multiple comparisons). For NB and WT, each treatment included 10 biological replicates, and the same root elongation data are also presented in Figure 2B. For the transposon mutants, each treatment included three biological replicates. Each biological replicate comprised two technical replicate plates with eight seedlings per plate. **(E)** Representative root growth inhibition (RGI) assay images. NB: no bacteria. WT: MF374 wild type. flg22-*Pa*: flg22 from *Pseudomonas aeruginosa*. Mock: sterile deionized water added as a control.

By screening 4,032 transposon mutants from the Tn5 library, we obtained five mutant candidates with a reduced ability to suppress GUS expression compared to the wild type (**Figure S2**). We mapped the transposon insertion sites in these candidate strains using arbitrary PCR and Sanger sequencing, identifying insertions in *ppiD*, *tonB1*, *BLR19_RS09010* (hereafter *RS09010*), *purD*, and *pmbA* (**Figure 1B, Table S2**). These genes encode proteins associated with protein folding and maturation (peptidylprolyl cis-trans isomerase D, PpiD) (*23*, *24*); TonB-dependent transport (TonB energy transducer, TonB1) (*25*, *26*); nucleic acid processing (3’-5’ exonuclease, encoded by *RS09010*) (*27*, *28*); purine biosynthesis (phosphoribosylamine-glycine ligase, PurD) (*29*); and proteolysis (metalloprotease component PmbA/TldE) (*30*).

We retested the five candidates in a normalized *pCYP71A12*::*GUS* reporter assay to determine the range of conditions under which each mutant retained immune attenuation activity. Because bacterial density and elicitor concentration both influence this phenotype, as we showed previously (*17*), we tested initial bacterial densities ranging from OD_600_ =10^-2^ to 10^-4^ with flg22-*Pa* at 1 or 0.1 µM. GUS activation, as indicated by blue staining of the root tip, was scored across approximately 10 to 17 seedlings per condition. Wild type MF374 prevented flg22-*Pa*-triggered GUS activation at all bacterial and flg22 concentrations tested, while the candidate mutants showed different degrees of loss of this phenotype (**Figure 1C**). The *tonB1*::Tn mutant showed the strongest defect, retaining immune attenuation only under the least stringent conditions tested: lower flg22-*Pa* concentration (0.1 µM) and higher bacterial density (OD_600_ = 10^-2^ and 10^-3^). The *ppiD*::Tn mutant showed a similar but slightly less severe defect to the *tonB1*::Tn strain, additionally retaining immune attenuation at the highest bacterial loading (OD_600_ = 10^-2^) with 1 µM flg22-*Pa*. The *purD*::Tn mutant showed the weakest defect, with a loss of attenuation only at 1 µM flg22-*Pa* and OD_600_ 10^-4^. The *RS09010*::Tn and *pmbA*::Tn mutants attenuated GUS activation at all conditions tested, showing no loss of activity relative to the wild type.

To validate these candidates using an independent immune activation-associated phenotype, we tested the five Tn mutants in a root growth inhibition (RGI) assay using Arabidopsis *pWER*::*FLS2-GFP* seedlings treated with 1 µM flg22-*Pa* (**Figure 1D,E**). In the absence of bacteria, root growth was strongly inhibited by flg22-*Pa*-triggered immune activation in this hypersensitive plant line. MF374 WT significantly suppressed the flg22-*Pa*-triggered RGI. Among the five mutant candidates, *ppiD*::Tn and *tonB1*::Tn showed a severe RGI phenotype under flg22-*Pa* treatment, with root lengths approaching those observed in the no bacteria control. The other mutants, *purD*::Tn, *RS09010*::Tn, and *pmbA*::Tn, suppressed RGI in the presence of flg22-*Pa* similarly to the wild type. Together, the GUS and RGI assays identified *tonB1* and *ppiD* as genes required for the robust attenuation of flg22-*Pa*-triggered immune responses by MF374.

### Targeted gene deletions validate MF374 loci contributing to immune attenuation

Because transposon insertions can incompletely disrupt gene function and can also have polar effects on neighboring genes, we constructed targeted gene deletions of *ppiD* and *tonB1* to validate their roles in attenuation of flg22-triggered immune activation. The *ppiD* gene is located at the beginning of a locus containing *trpEXGDC*, including genes annotated for tryptophan biosynthesis (*trpE*, *trpG*, *trpD*, and *trpC*) as well as a *trpX*, a gene oriented in the opposite direction that encodes a small hypothetical protein (**Figure 1B**). In other Alphaproteobacteria, *ppiD* genes are reported to be co-located and co-regulated with *trp* biosynthesis genes, including the *ppiD-trpDC-moaC-moeA* operon in *Sinorhizobium meliloti* (*31*). In MF374, *ppiD* and *trpE* overlap by four bases, raising the possibility that the *ppiD*::Tn phenotypes could reflect disruption of *ppiD,* polar effects on the neighboring *trpEXGDC* locus, or both. To distinguish these possibilities, we constructed markerless deletions Δ*ppiD* and Δ*trpEXGDC* (**Table 1**). A markerless *tonB1* deletion could not be recovered, suggesting strong selective pressure to retain this gene; therefore, *tonB1* was disrupted with a gentamicin resistance cassette, resulting in a Δ*tonB1*::*Gm^R^*strain.

**Table 1.** List of *Brevundimonas* sp. MF374 gene knockouts and gene complements.

| Strain ID | Strain w/ vector | Genotype |
| --- | --- | --- |
| MF374_CL001 | MF374 w/ pCL001 | $\Delta ppiD$ |
| MF374_CL002 | MF374 w/ pCL002 | $\Delta ppiD$-trpEXGDC |
| MF374_CL003 | MF374 w/ pCL012 | $\Delta tonB1::Gm^R$ |
| MF374_CL004 | MF374 w/ pCL015 | $\Delta trpEXGDC$ |
| MF374_CL005 | MF374_CL003 w/<br>pCL001 | $\Delta tonB1::Gm^R \Delta ppiD$ |
| MF374_CL006 | MF374_CL003 w/<br>pCL015 | $\Delta tonB1::Gm^R \Delta trpEXGDC$ |
| MF374_CL007 | MF374_CL001 w/<br>pCL007 | $\Delta ppiD$ / pBBR1::ppiD |
| MF374_CL008 | MF374_CL002 w/<br>pCL007 | $\Delta ppiD$-trpEXGDC /<br>pBBR1::ppiD |
| MF374_CL009 | MF374_CL002 w/<br>pCL008 | $\Delta ppiD$-trpEXGDC /<br>pBBR1::trpEXGDC |
| MF374_CL010 | MF374_CL002 w/<br>pCL013 | $\Delta ppiD$-trpEXGDC /<br>pBBR1::ppiD-trpEXGDC |
| MF374_CL011 | MF374_CL003 w/<br>pCL009 | $\Delta tonB1::Gm^R$ / pBBR1::tonB1 |
| MF374_CL012 | MF374_CL004 w/<br>pCL008 | $\Delta trpEXGDC$ /<br>pBBR1::trpEXGDC |

\*X in trpEXGDC represents a hypothetical gene (*BLR19\_RS02890*).

We tested the Δ*tonB1*::*Gm^R^,* Δ*ppiD,* and Δ*trpEXGDC* deletion mutants using the normalized GUS assay across a bacterial dilution series from OD_600_ = 1 to 10^-5^ with 1 µM flg22-*Pa* (**Figure 2A**). MF374 WT attenuated flg22-*Pa*-triggered immune activation from OD_600_ = 1 to 10^-4^. This attenuation was reduced in all three mutants. In Δ*tonB1*::*Gm^R^*, attenuation was eliminated from OD_600_ = 10^-2^ to 10^-4^, remaining only at the highest bacterial loadings. Δ*ppiD* and Δ*trpEXGDC* showed weaker defects, with loss of attenuation observed at OD_600_ = 10^-4^.

**Figure 2.**
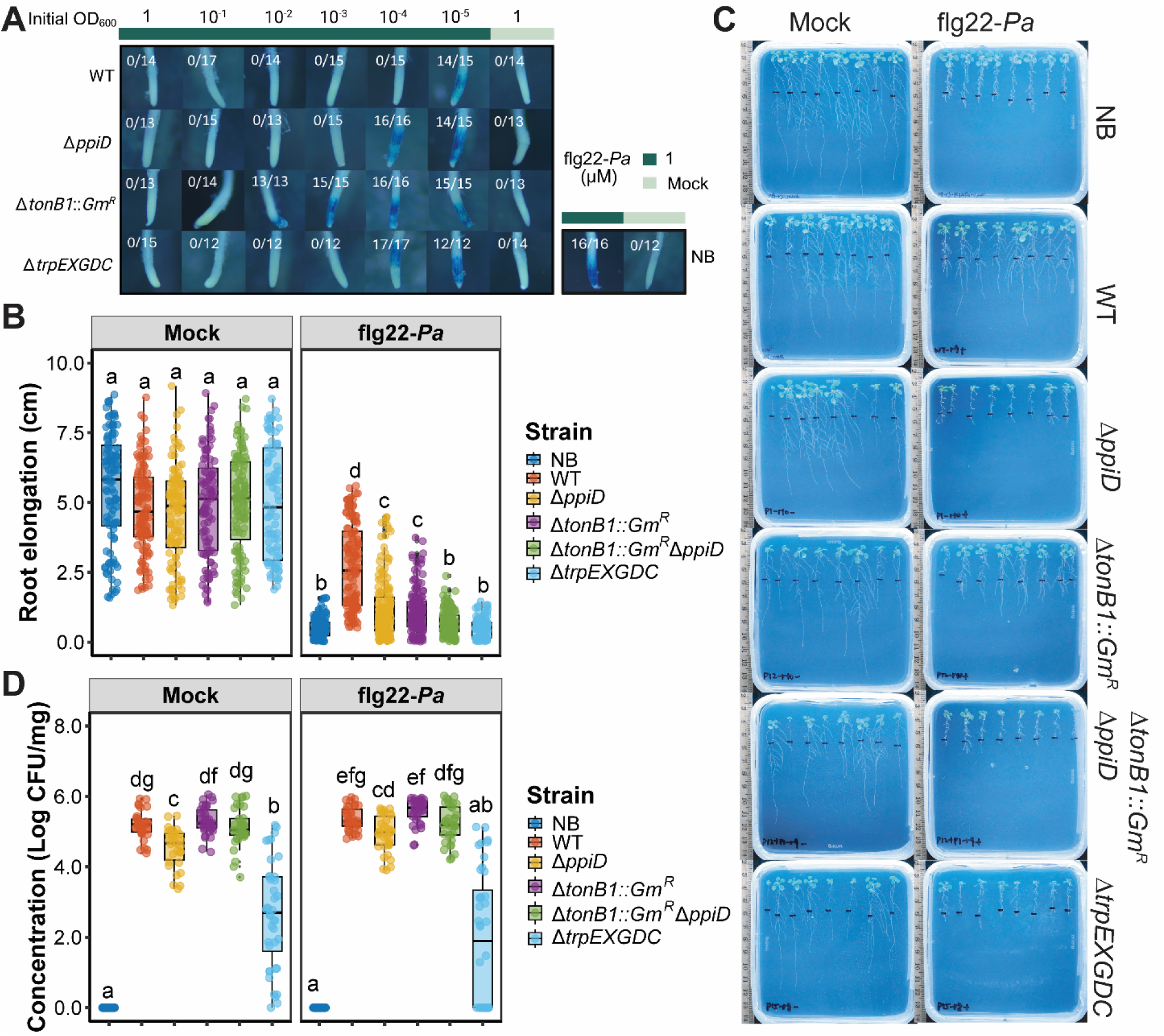
Disruption of *tonB1* impairs immune attenuation by *Brevundimonas* sp. MF374 without reducing root colonization. **(A)** GUS reporter assays of MF374 wild type (WT) and deletion mutants in *A. thaliana pCYP71A12*::*GUS* at initial OD_600_ of 1 to 10^-5^ with 1 µM flg22-*Pa*. Fractions indicate the number of blue-stained root tips relative to the total number examined across three biological replicates. **(B)** Root elongation of *A. thaliana pWER::FLS2-GFP* seedlings 9 days after inoculation with no bacteria (NB), wild type (WT), or deletion mutant strains at OD_600_ = 0.01, with or without 1 µM flg22-*Pa*. Each treatment included 10 biological replicates, each comprising two technical replicate plates with eight seedlings per plate. The same NB and WT root elongation data are also presented in Figure 1D. **(C)** Representative root growth inhibition (RGI) assay images. **(D)** Root colonization of *A. thaliana pWER*::*FLS2-GFP* by WT and mutant strains 9 days after inoculation, expressed as Log (CFU per mg root tissue). Colonization was quantified from 10 biological replicates with two technical replicates from the experiments in (B). Different letters across mock and flg22-*Pa* panels in (B) and (D) indicate significant differences (*P* < 0.05; Kruskal-Wallis test followed by Dunn’s test with the Benjamini-Hochberg method for *P*-value adjustment of multiple comparisons). NB: no bacteria. WT: MF374 wild type. flg22-*Pa*: flg22 from *Pseudomonas aeruginosa*. Mock: sterile deionized water control.

Because deletion of all three loci disrupted the attenuation of GUS activation to some extent, we next constructed pairwise double deletion mutants to determine whether combining these gene deletions produced stronger attenuation defects than the single deletions. This resulted in the construction of Δ*tonB1*::*Gm^R^* Δ*ppiD*, Δ*ppiD*-*trpEXGDC*, and Δ*tonB1*::*Gm^R^* Δ*trpEXGDC* mutant strains (**Table 1**). We tested these double mutants using the same normalized GUS assay (**Figure S3**). The Δ*tonB1*::*Gm^R^* Δ*ppiD* mutant did not show a stronger attenuation defect than the Δ*tonB1*::*Gm^R^* alone (**Figure S3**). Similarly, the Δ*tonB1*::*Gm^R^* Δ*trpEXGDC* resembled Δ*tonB1*::*Gm^R^* (**Figure S3**). These results suggest that *tonB1* is the primary determinant of this loss of attenuation phenotype, with no additional defect observed with the addition of deletion of *ppiD* or the *trp* locus. By contrast, the Δ*ppiD*-*trpEXGDC* mutant showed a stronger attenuation defect than either the Δ*ppiD* or Δ*trpEXGDC* deletion alone, with loss of attenuation observed at OD_600_ = 10^-3^ (**Figure S3**). To determine whether the loss of attenuation phenotypes could be rescued, we introduced complementation constructs driven by a *lacZ* promoter on the broad host range plasmid pBBR1-MCS2 into the corresponding mutant backgrounds. These constructs included pBBR1::*ppiD,* pBBR1::*tonB1,* pBBR1::*trpEXGDC,* and pBBR1::*ppiD*-*trpEXGDC* with strains summarized in **Table 1**. We tested complementation in relevant single and double deletion strains using the same normalized GUS assay (**Figure S4**). In this GUS assay, wild type MF374 attenuated GUS activation down to OD_600_ = 10^-4^ (Figure 2A), providing a benchmark for full restoration of the wild type phenotype. Complementation of Δ*tonB1*::*Gm^R^* with pBBR1::*tonB1* restored attenuation at OD_600_ = 10^-2^ and 10^-3^ compared to Δ*tonB1*::*Gm^R^* (**Figure S4A and Figure 2A**). Complementation of Δ*trpEXGDC* with pBBR1::*trpEXGDC* restored the attenuation to OD_600_ = 10^-4^, matching the WT (**Figure S4A and Figure 2A**). In the Δ*ppiD-trpEXGDC* background, complementation with pBBR1::*ppiD*, pBBR1::*trpEXGDC*, or the full pBBR1::*ppiD-trpEXGDC* construct gave similar results, with partial restoration of attenuation observed at OD_600_ = 10^-3^ (**Figure S4B and Figure S3**). By comparison, pBBR1::*ppiD* did not restore attenuation in the Δ*ppiD* single mutant background (**Figure S4A and Figure 2A**). Overall, complementation restored the attenuation phenotype in several mutant backgrounds, although in most cases not to the level of the MF374 wild type. This incomplete complementation may reflect non-native or suboptimal expression levels of the genes from the plasmid-based constructs in this non-model bacterium.

To determine whether these GUS phenotypes extended to RGI, we tested the targeted mutants in the *pWER*::*FLS2-GFP* RGI assay using 1 µM flg22-*Pa* and bacteria inoculated at OD_600_ = 10^-2^ (**Figure 2B,C**). MF374 WT significantly suppressed 1 µM flg22-*Pa*-triggered RGI. The Δ*ppiD*, Δ*tonB1*::*Gm^R^*, and Δ*trpEXGDC* single mutants and Δ*tonB1*::*Gm^R^* Δ*ppiD* double mutant all showed significant reduction in RGI suppression. The strongest reductions in root elongation were observed for the Δ*trpEXGDC* and Δ*tonB1*::*Gm^R^*Δ*ppiD* double mutant (**Figure 2B**). Because RGI suppression could be influenced by bacterial colonization, we quantified root colonization by each strain under mock and flg22-*Pa*-treated conditions (**Figure 2D**). These results showed that Δ*ppiD* and Δ*trpEXGDC* had significantly lower colonization (Log CFU/mg root fresh weight, *P* < 0.05) than MF374 WT, with the Δ*trpEXGDC* colonization defect being more severe than Δ*ppiD* (**Figure 2D**). Interestingly, flg22-*Pa* treatment did not significantly affect colonization of any strain compared to its colonization pattern without flg22, indicating that flg22-*Pa*-triggered immune activation did not measurably reduce MF374 colonization under these conditions even in mutants with severe defects in attenuation. This was most evident for Δ*tonB1*::*Gm^R^*, which showed a strong loss of immune attenuation but colonized roots similarly to WT MF374.

This substantial colonization defect suggested that the poor RGI suppression observed for Δ*trpEXGDC* (**Figure 2B,C**) could reflect reduced bacterial fitness rather than a direct defect in immune attenuation. We therefore examined the effects of tryptophan availability on bacterial growth and immune attenuation (**Figure S5**). In MS medium supplemented with 1% Casamino acids, which lacks tryptophan, Δ*trpEXGDC* showed no detectable growth, whereas growth was restored by tryptophan supplementation or complementation with *trpEXGDC* (**Figure S5A,B**). The original *ppiD::Tn* mutant similarly showed a strong tryptophan-responsive growth defect, consistent with a possible polar effect on the overlapping downstream *trpEXGDC* locus, whereas the clean Δ*ppiD* mutant retained growth without added tryptophan (**Figure S5A–D**). By contrast, disruption of *tonB1* did not cause a comparable growth defect. Consistent with a contribution of tryptophan limitation to the Δ*trpEXGDC* phenotype, supplementation with 0.1mM tryptophan restored immune attenuation by Δ*trpEXGDC* in the GUS assay (**Figure S5E**).

Taken together, these assays identified *tonB1* as the strongest contributor to MF374-mediated attenuation of flg22-*Pa*-induced immunity, with Δ*tonB1*::*GmR* showing severe defects in both GUS and RGI assays without a corresponding loss of root colonization. The *ppiD-trpEXGDC* locus also affected attenuation-associated phenotypes, but these effects were weaker than *tonB1* in the GUS assay and were complicated by colonization defects in the RGI assay. In particular, the severe loss of RGI suppression by Δ*trpEXGDC* may reflect reduced plant-associated fitness rather than a direct defect in immune attenuation. Ultimately, because these genetic results showed that each locus affected at least one attenuation phenotype, we carried these mutants forward to test what interactions among MF374, flg22-*Pa*, and the plant are required for immune attenuation.

### Intact MF374 cells deplete flg22 immune-eliciting activity from the extracellular environment

To determine how MF374 attenuates flg22-triggered immunity, we developed a panel of five GUS assay variants (A1-A5, **Figure 3A**) to separate the contributions of bacterial cells, plant-associated bacteria, and cell-free components in the attenuation of flg22-induced GUS expression. The standard normalized GUS assay served as the reference assay condition (A1). To distinguish whether MF374 attenuated subsequent immune activation by modifying the plant during the initial bacteria-seedling incubation or through activity present in the medium when flg22 was subsequently added, we separated these components after 14 h. The unfiltered medium containing suspended MF374 cells was transferred to new reporter seedlings before addition of flg22-*Pa* in the second assay variant (A2), whereas the original MF374-exposed seedlings, including any colonized bacteria, were washed and transferred to fresh medium before flg22-*Pa* treatment in the third assay variant (A3). To further determine whether attenuation in the medium resulted from extracellular factors produced before elicitor exposure or required interaction of flg22-*Pa* with intact MF374 cells, we compared two additional conditions. In A4, the medium from the initial MF374-seedling incubation was filtered to remove bacterial cells, transferred to new reporter seedlings, and then supplemented with flg22-*Pa*. In A5, flg22-*Pa* was first incubated with the bacteria-containing medium for 5 h, after which MF374 cells were removed by filtration and the resulting cell-free medium was transferred to new reporter seedlings.

**Figure 3.**
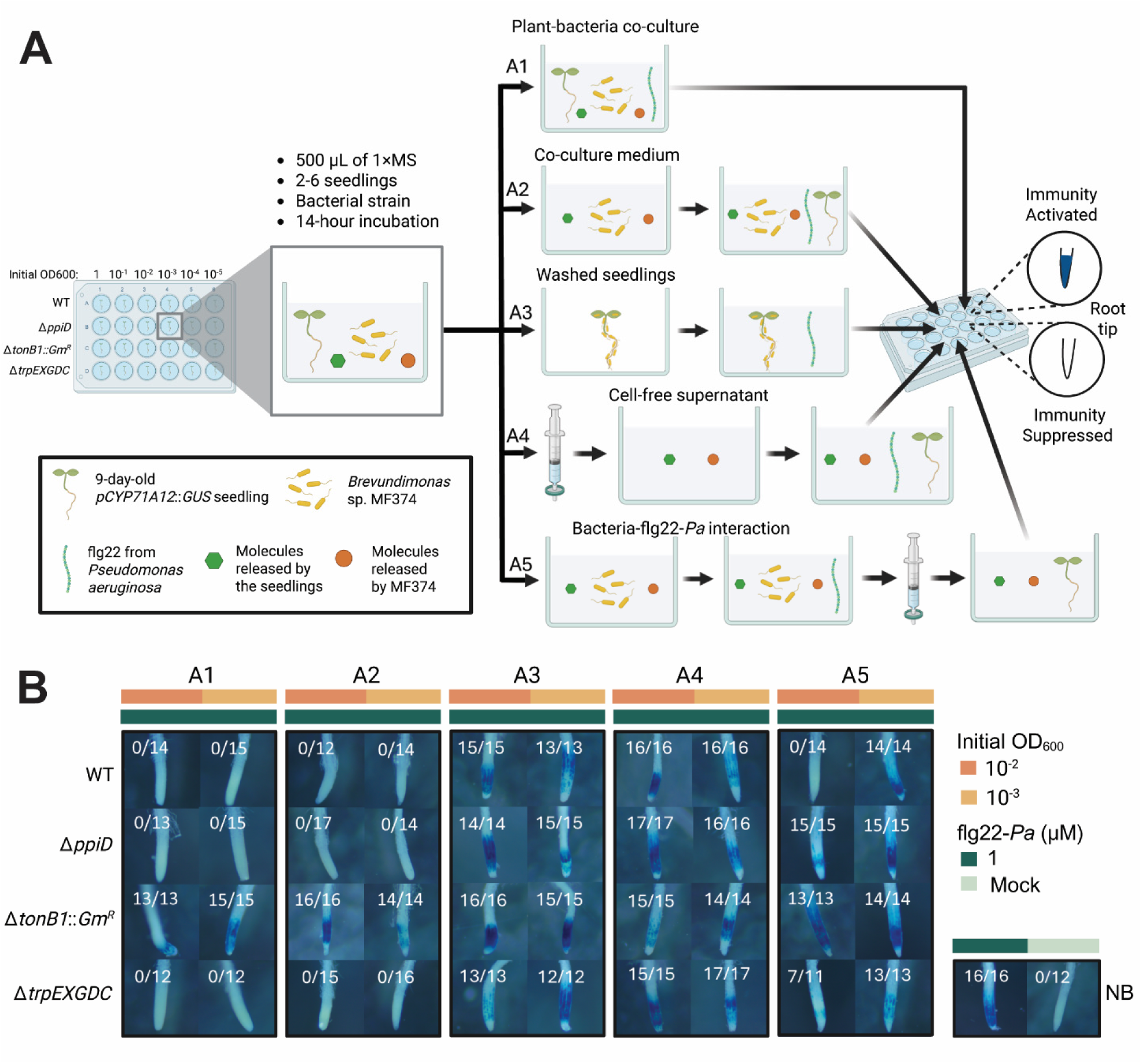
Direct interaction with *Brevundimonas* sp. MF374 cells reduces the immune-eliciting activity of flg22-*Pa* in a TonB1-dependent manner. **(A)** Schematic of five GUS reporter assays designed to distinguish the contributions of bacterial cells, secreted factors, and plant-associated effects to immune attenuation. Assays included the standard GUS assay plant-bacteria co-culture (A1), transfer of co-culture medium containing bacteria to fresh seedlings (A2), washed seedlings after co-culture transferred to fresh medium (A3), cell-free filtered co-culture supernatant (A4), and flg22-*Pa* pre-incubated with bacteria followed by removal of bacterial cells by filtration (A5). Created with BioRender.com **(B)** GUS reporter assays of MF374 wild type (WT) and deletion mutants using *A. thaliana pCYP71A12*::*GUS* seedlings, with bacteria at initial OD_600_ values of 10⁻² or 10⁻³ and 1 µM flg22-*Pa*. Fractions indicate the number of blue-stained root tips relative to the total number examined across three biological replicates. Additional results for assays A2–A5 across the full bacterial dilution series are shown in Fig. S6. Images at OD₆₀₀ = 10⁻² and 10⁻³ are shared between Fig. 3B and Fig. S6, while the A1 images are also presented in Fig. 2A. NB, no bacteria; mock, sterile deionized water control.

The GUS assay panel showed that robust attenuation activity tracked with the presence of intact MF374 cells (**Figure 3B, Figure S6A-D**). Wild type MF374 attenuated flg22-*Pa*-induced GUS expression in both the standard assay (A1) and the assay where bacteria-containing medium is transferred to new seedlings (A2), both at bacterial loading of OD_600_ = 10^-2^ and 10^-3^. In contrast, washed MF374-exposed seedlings transferred to fresh medium (A3) did not attenuate flg22-*Pa*-induced GUS expression. Cell-free medium generated by filtering out MF374 cells before flg22-*Pa* addition (A4) also failed to attenuate GUS expression at these bacterial densities. Together, these results suggest that robust attenuation required the presence of intact MF374 cells at the time of elicitor exposure. To test whether interaction with intact MF374 cells directly altered flg22-*Pa* activity, we next examined A5, in which flg22-*Pa* was incubated with MF374 before the bacterial cells were removed. In A5, interaction with MF374 cells at OD_600_ = 10^-2^ abolished the immune-eliciting activity of flg22-*Pa*, although this was not the case at the lower bacterial loading of OD_600_ = 10^-3^. Thus, interaction with intact MF374 cells was sufficient to deplete flg22-*Pa* immune-eliciting activity from the extracellular environment.

The targeted mutants further distinguished the genetic requirements for this cell-dependent depletion of flg22-*Pa* activity. Across the bacterial dilution series, the mutant phenotypes in the bacteria-containing conditions (A1 and A2) broadly recapitulated the preceding results (**Figure 2A**), where Δ*ppiD* and Δ*trpEXGDC* showed modest reductions in attenuation relative to wild type, and Δ*tonB1*::*Gm^R^* showed a substantially stronger defect (**Figure 3B; Figure S6A**). None of the mutants attenuated flg22-*Pa*-induced GUS expression in the washed-seedling condition (A3) and cell-free supernatants condition (A4) (**Figure 3B, Figure S6B,C**). The A5 assay provided the clearest test of which genes contribute to attenuation mediated by interaction of flg22-*Pa* with intact MF374 cells. At OD_600_ = 10^-2^, both Δ*tonB1*::*Gm^R^* and Δ*ppiD* failed to abolish the immune-eliciting activity of flg22-*Pa*, whereas Δ*trpEXGDC* showed a partial phenotype with GUS activation in 7 of 11 seedlings (**Figure 3B**). Across the full dilution series (**Figure S6D**), wild type MF374 abolished flg22-*Pa* activity following incubation at bacterial densities down to OD_600_ = 10^-2^, whereas Δ*ppiD* required a higher bacterial density (OD_600_ = 10^-1^) to retain this phenotype. The Δ*tonB1*::*Gm^R^* strain again showed the strongest defect, with comparable attenuation observed only at the highest bacterial loading tested (**Figure S6D**). This could be consistent with a role for TonB-dependent uptake in reducing extracellular flg22-*Pa* activity. Together, these results indicate that efficient depletion of flg22-*Pa* immune-eliciting activity requires interaction with intact MF374 cells and is strongly dependent on *tonB*, with *ppiD* also contributing to this process.

### TonB contributes to MF374-mediated attenuation of multiple immune eliciting peptides

We next asked whether MF374-mediated immune attenuation extended beyond flg22-*Pa* to other immune elicitors. We tested a distinct flg22 sequence from *Dyella japonica* MF79 (flg22-*Dja*), AtPEP1, elf18, and chitin. Flg22-*Dja* differs in sequence from flg22-*Pa* and has previously been shown to elicit Arabidopsis immune responses at levels slightly weaker than flg22-*Pa*, including in studies of MF79-mediated immune suppression (*3*, *17*). Of these elicitors, only flg22-*Dja* activated the *pCYP71A12*::*GUS* reporter under the conditions tested (**Figure S7A**). We therefore examined flg22-*Dja* using the A1, A4, and A5 assay conditions developed for flg22-*Pa*. Wild type MF374 attenuated flg22-*Dja*-induced GUS expression in the standard assay (A1), while cell-free supernatant generated before elicitor addition (A4) showed little attenuation at the bacterial densities examined. As with flg22-*Pa*, interaction of flg22-*Dja* with intact MF374 cells (A5) substantially reduced its subsequent immune-eliciting activity (**Figure 4A, Figure S8**). Consistent with the flg22-*Pa* results, Δ*tonB1*::*GmR* again showed the strongest defect in attenuation with flg22-*Dja*, whereas Δ*ppiD* and Δ*trpEXGDC* showed weaker phenotypes (**Figure 4A, Figure S8**).

**Figure 4.**
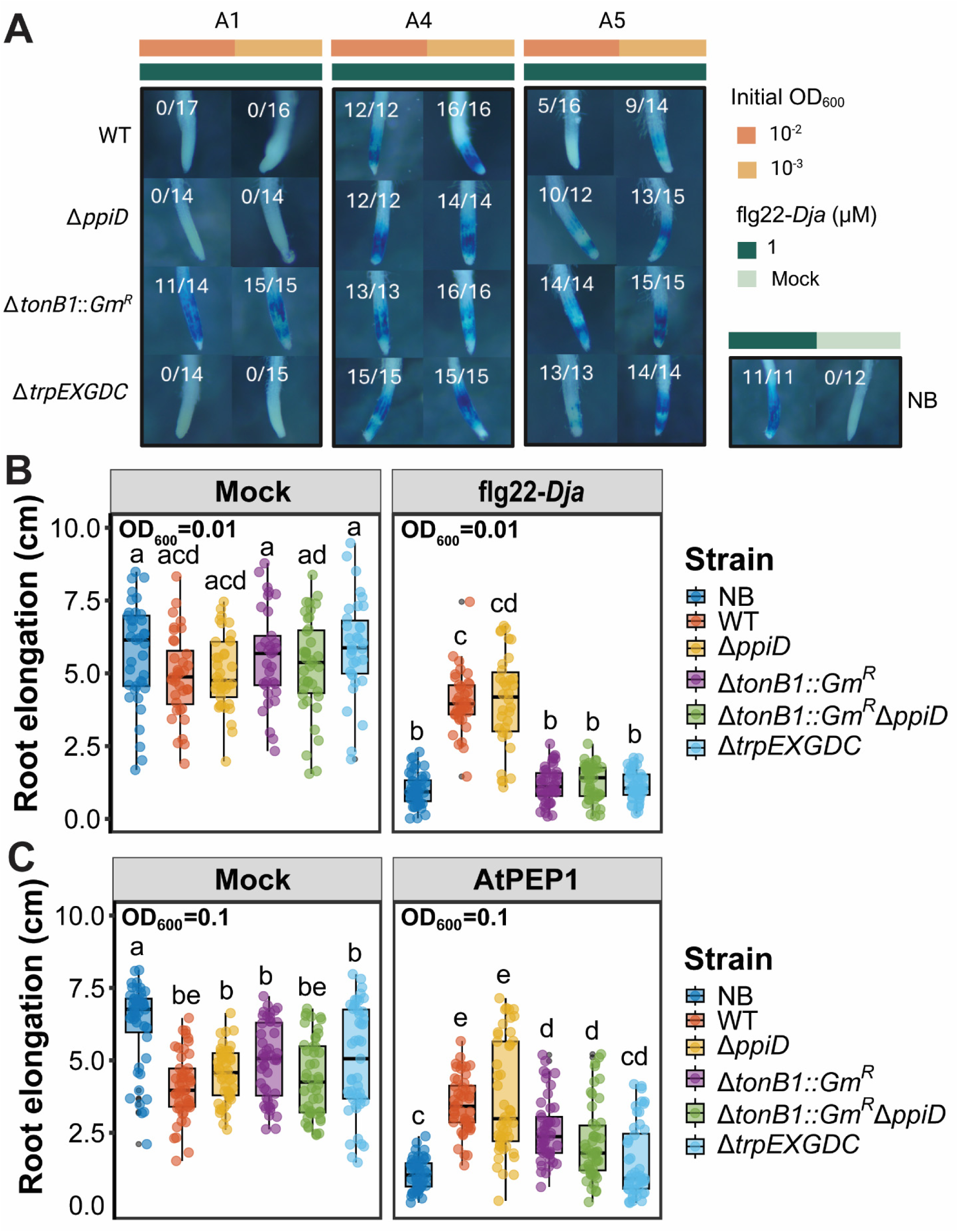
TonB1 contributes to MF374-mediated attenuation of plant immune responses triggered by flg22-*Dja* and AtPEP1. **(A)** GUS reporter assays of MF374 wild type (WT) and deletion mutants using *A. thaliana pCYP71A12*::*GUS* seedlings to evaluate flg22-*Dja*-triggered immunity in plant bacteria co-cultures (A1), cell-free supernatant (A4), and following pre-incubation of flg22-*Dja* with bacterial cells (A5). Bacteria were applied at OD_600_ = 10^-2^ and 10^-3^ with 1 μM flg22-*Dja*. Fractions indicate the number of blue-stained root tips relative to the total number examined across three biological replicates. Additional results across the full bacterial dilution series are shown in Fig. S8. **(B)** Root elongation of *A. thaliana pWER::FLS2-GFP* seedlings 9 days after inoculation with no bacteria (NB), wild type (WT), or deletion mutant strains (OD_600_ = 0.01), with or without 1 µM flg22-*Dja*. **(C)** Root elongation of *A. thaliana* Col-0 seedlings 9 days after inoculation with no bacteria (NB), wild type (WT), or deletion mutant strains (OD_600_ = 0.1), with or without 0.1 µM AtPEP1. For (B) and (C), each treatment included three biological replicates, each comprising two technical replicate plates with approximately eight seedlings per plate. Different letters across both mock and flg22-*Dja* panels in **(B)** or both mock and AtPEP1 panels in **(C)** indicate significant differences (*P* < 0.05; Kruskal-Wallis test followed by Dunn’s test with the Benjamini-Hochberg method for *P*-value adjustment of multiple comparisons). flg22-*Dja*: flg22 peptide from *Dyella japonica* MF79; AtPEP1: Arabidopsis thaliana Plant Elicitor Peptide 1; NB: no bacteria; mock: sterile deionized water control.

Because AtPEP1, elf18, and chitin did not activate the *pCYP71A12*::*GUS* reporter, we evaluated these elicitors using the RGI assays in *A. thaliana* Col-0. Consistent with previous observations of AtPEP1-induced root growth inhibition and its attenuation by root commensals (*19*), AtPEP1 strongly reduced root elongation under our assay conditions, and this inhibition was significantly attenuated by MF374 across the bacterial densities tested (**Figure 4C, Figure S7B**). In contrast, significant attenuation of elf18- or chitin-induced root growth inhibition was not observed (**Figure S7B**). Because wild type MF374 attenuated both flg22-*Dja*- and AtPEP1-induced root growth inhibition, we next asked whether the genes identified for flg22-*Pa* attenuation also contributed to these phenotypes (**Figure 4B,C**). For flg22-*Dja*, mutants containing the *tonB1* disruption showed strong defects in attenuation, whereas Δ*ppiD* retained activity comparable to wild type (**Figure 4B**). A similar pattern was observed with AtPEP1, where disruption of *tonB1*, either alone or together with *ppiD*, strongly reduced MF374-mediated attenuation, whereas Δ*ppiD* retained attenuation comparable to wild type (**Figure 4C**). Δ*trpEXGDC* also showed reduced attenuation in both assays, although interpretation of this phenotype is complicated by the growth and colonization defects associated with this mutant (**Figure 2D, Figure S5**). Together, these results show that the contribution of *tonB* to MF374-mediated immune attenuation extends beyond flg22-*Pa* to both flg22-*Dja* and AtPEP1.

## Discussion

*Brevundimonas* sp. MF374 has previously emerged as a strong suppressor of Arabidopsis flg22-triggered immunity (*17*, *19*). Despite this strong phenotype, the bacterial mechanisms responsible for immune suppression by MF374 remained unknown. Here, we generated an RB-TnSeq library in MF374 and used a forward genetic screen to identify bacterial genes required for this robust immune attenuation (**Figure 1**). Subsequent targeted mutant analysis identified *tonB1* as the strongest determinant of MF374-mediated immune attenuation, while disruption of *ppiD* caused a weaker but reproducible loss of attenuation (**Figure 2**). Using a series of modified GUS assays designed to separate the contributions of bacterial cells, exposed seedlings, and cell-free medium, we further examined how MF374 reduced flg22-triggered immune activation (**Figure 3**). Interaction of flg22 with intact MF374 cells prior to seedling treatment substantially reduced its ability to elicit an immune response on reporter seedlings, and this phenotype was strongly dependent on *tonB1*. The contribution of *tonB1* was also observed in the attenuation of immune activation by a distinct flg22 sequence, flg22-*Dja*, and AtPEP1 (**Figure 4**). These results identify a TonB1-dependent process as a major component of immune attenuation by MF374 and support a model in which MF374 reduces the extracellular availability of immune-eliciting peptides.

The functions of TonB1 and PpiD suggest a model (**Figure 5**) in which transport and protein processing functions contribute to depletion of extracellular immune elicitors by MF374. TonB1 functions as part of TonB1-ExbB-ExbD, which couples the proton motive force across the inner membrane to TonB-dependent transporters (TBDTs) in the outer membrane to power uptake of extracellular substrates (*25*, *32*). The strong requirement for *tonB1* in the intact-cell depletion assay (A5, **Figure 3**) is therefore consistent with TonB1-dependent transport reducing the extracellular availability of flg22. PpiDs are inner membrane-associated chaperones that participate in the early folding and maturation of newly translocated proteins (*23*, *33*, *34*), including inner membrane proteins (IMPs) and outer membrane proteins (OMPs) (*33*, *35*). PpiDs also assist with releasing OMPs into the periplasm (*24*, *36*, *37*), where they can subsequently be engaged by chaperones such as SurA and delivered to the β-barrel assembly machinery (BAM) complex for folding and insertion into the outer membrane (*37–40*). Because TonB1 is inner membrane anchored and TBDTs are outer membrane proteins, these functions of PpiDs provide a potential route by which PpiD in MF374 could influence the assembly or activity of the TonB1 transport machinery involved in immune elicitor depletion.

**Figure 5.**
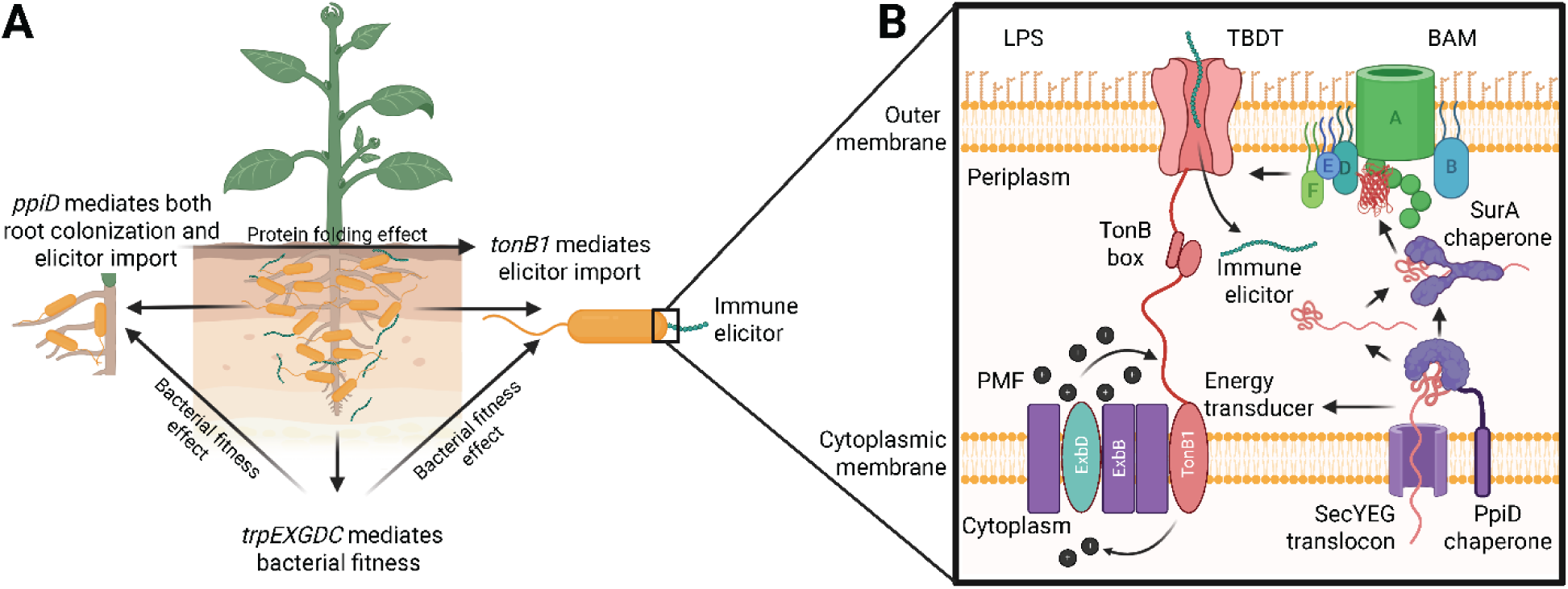
Proposed model of TonB1-dependent immune attenuation by *Brevundimonas* sp. MF374. **(A)** Summary of the contributions of *tonB1*, *ppiD*, and *trpEXGDC* to immune attenuation. TonB1 and PpiD contribute to the reduction of immune-eliciting activity, while PpiD also contributes to root colonization. The *trpEXGDC* locus is required for tryptophan biosynthesis, bacterial fitness, and root colonization. **(B)** Proposed mechanism of TonB-dependent immune elicitor uptake. The TonB-ExbB-ExbD complex uses the proton motive force (PMF) to energize outer membrane TonB-dependent transporters (TBDTs), potentially enabling the import of immune elicitors. PpiD may facilitate maturation of TBDTs and other outer membrane proteins through interactions with the SecYEG translocon and subsequent protein folding pathways involving periplasmic chaperone SurA and β-barrel assembly machinery (BAM) complex (BamABDEF). LPS: lipopolysaccharide. Created with BioRender.com

TonB-dependent transport in immune attenuation by plant-associated commensals has precedent in *Rhodanobacter* sp. Root179 (hereafter, R179). Ordon *et al.* generated a transposon mutant library in R179 and screened approximately 7,100 mutants for impaired suppression of flg22-induced root growth inhibition, identifying *dssB*, which encodes a TonB-dependent transporter, as one of the major genetic determinants of this phenotype (*21*). Targeted deletion of *dssB* impaired suppression of flg22-induced root growth inhibition and immune-associated *FRK1* expression without substantially affecting bacterial growth or root colonization (*21*). However, the role of DssB in this phenotype was not linked to direct depletion or transport of flg22. To place the TonB-dependent phenotypes observed in MF374 and R179 in broader context, we compared the TonB and TBDT repertoires of these strains with two other well-characterized Gram-negative plant-associated immune suppressors, *Dyella japonica* MF79 and *Pseudomonas simiae* WCS417, together with *E. coli* K-12 as a reference strain (**Figure S9; Table S10**). These strains encode diverse repertoires of TonB proteins and TBDTs, including four TonB proteins and 35 predicted TBDTs in MF374 (**Figure S9; Table S10**). Phylogenetic comparison did not reveal a close relationship between the MF374 TonB1 identified in this study and the TonB proteins encoded by R179 (**Figure S9A**), nor did MF374 contain an obvious close homolog of the R179 DssB TBDT (**Figure S9B**). Thus, although TonB-dependent transport contributes to immune attenuation in both MF374 and R179, comparisons of the relevant transport components identified through genetic screening do not point to a single conserved TonB/TBDT pair underlying these immune attenuation phenotypes.

We also examined the contribution of the *trpEXGDC* locus in immune attenuation because it lies immediately downstream of *ppiD*, overlapping with the *ppiD* coding sequence. This raised the possibility that the original *ppiD*::Tn insertion affected expression of the downstream tryptophan biosynthesis genes. A similar genomic association between *ppiD* and tryptophan biosynthesis genes has been reported in other Alphaproteobacteria, including *Sinorhizobium meliloti* (*31*). Targeted disruption of *trpEXGDC* produced defects in immune attenuation in both GUS and RGI assays, but this mutant also showed a severe defect in root colonization (**Figure 2D**) and no detectable growth in medium lacking exogenous tryptophan (**Figure S5A**). Growth was restored by tryptophan supplementation or genetic complementation (**Figure S5B**), and addition of 0.1 mM tryptophan also restored attenuation of flg22-induced GUS activation (**Figure S5E**). These results indicate that the apparent contribution of *trpEXGDC* to immune attenuation is largely associated with its requirement for bacterial growth and fitness rather than a specific role in TonB-dependent depletion of flg22. Consistent with this interpretation, the original *ppiD*::Tn mutant also showed a strong tryptophan-responsive growth defect, whereas the clean *ppiD* deletion strain, which leaves the *trpE* start codon intact, retained growth in the absence of added tryptophan. Thus, the *ppiD* and *trpEXGDC* phenotypes can be separated, supporting a role for PpiD in immune attenuation independent of the downstream tryptophan biosynthesis locus.

The contribution of *tonB1* to immune attenuation extended beyond flg22-*Pa*. MF374 also reduced the immune-eliciting activity of the distinct flg22 sequence flg22-*Dja*, and disruption of *tonB1* again produced the strongest defect in this attenuation (**Figure 4A,B, Figure S8**). Disruption of *ppiD* also reduced attenuation of flg22-*Dja* immune activation in the GUS assay, as with flg22-*Pa* (**Figure 4A, Figure S8**). Because AtPEP1, elf18, and chitin did not activate the *pCYP71A12::GUS* reporter under the conditions tested (**Figure S7A**), we evaluated these elicitors using the RGI assay. MF374 significantly attenuated AtPEP1-induced root growth inhibition in a *tonB*-dependent manner, with no contribution observed for *ppiD* (**Figure 4C, Figure S7B**). In contrast, MF374 did not significantly attenuate root growth inhibition triggered by elf18 or chitin under the conditions tested (**Figure S7B**). Together, these results establish TonB1 as a broader determinant of MF374-mediated immune attenuation, extending from distinct flg22 peptides to the endogenous plant elicitor AtPEP1.

We also show that interaction of flg22 with intact MF374 cells is required for the immune attenuation phenotype. This requirement distinguishes the MF374 mechanism from suppression mediated by extracellular factors, such as the secreted protease IssA produced by *D. japonica* MF79, which maintains strong suppressive activity in the cell-free supernatant (*17*). Together with the strong genetic requirement for *tonB1*, the results presented here support TonB1-dependent transport as the most likely mechanism for depletion of active extracellular flg22 by MF374. Additional or redundant cell-associated mechanisms could also contribute to the attenuation phenotype. For example, the requirement for intact cells could also permit contributions from surface-associated degradation or sequestration of flg22 at the bacterial cell surface. These additional mechanisms may explain attenuation in the Δ*tonB1* mutant observed at high bacterial loadings (**Figure S6**, **Figure S8**). The weaker contribution of *ppiD* is also consistent with an accessory role in this process, potentially by promoting proper folding or assembly of membrane or periplasmic proteins required for efficient immune elicitor depletion. Thus, while the specific TBDT and subsequent fate of flg22 remain to be determined, the genetic and functional data defines a cell-associated, TonB1-dependent mechanism that efficiently removes flg22 immune-eliciting activity from the extracellular environment.

Only a limited number of mechanisms used by root-associated commensals to attenuate plant immune activation have been characterized. *D. japonica* MF79 uses the secreted IssA proteases to degrade immune elicitors, while *Pseudomonas simiae* WCS417 modulates immune activation through changes in environmental pH (*17*, *18*, *20*). Genetic studies in *Rhodanobacter* sp. R179 further implicated a TonB-dependent transporter in attenuation of flg22-triggered immune responses, although how this transporter altered immune-elicitor activity remained unresolved (*21*). Here, we connect TonB1-dependent transport machinery to the cell-associated depletion of extracellular flg22 activity by *Brevundimonas* sp. MF374, defining a distinct mechanism by which a root-associated commensal can act directly on an immune elicitor before plant perception. The contribution of TonB1 to attenuation of distinct flg22 peptides and AtPEP1 further suggests that this strategy can influence responses to multiple immune elicitors. Together, these findings expand the known plant-microbe interaction mechanisms by which commensal root-associated bacteria attenuate plant immune activation.

## Materials and Methods

### Preparation of plants

In this study, the *Arabidopsis thaliana pCYP71A12*::*GUS* line was used in the β-glucuronidase (GUS) assay, and the *A. thaliana pWER*::*FLS2-GFP* and Col-0 (ecotype Columbia) lines were applied in the root growth inhibition (RGI) assay. The following protocol was adapted from a previous study (*19*) with some modifications. Fifty milligram seeds of each *A. thaliana* line were sterilized using 1.5 mL of washing solution containing 70% (v/v) household bleach (CLOROX) containing 8.25% (v/v) sodium hypochlorite, 29.8% (v/v) DI-water, and 0.2% (v/v) Tween-20 for 10 min on a vortex mixer (Fisherbrand) at 3,000 rpm and washed eight times with sterile DI-water. After stratification at 4 °C for 24 hours in the dark, the sterilized *A. thaliana pCYP71A12*::*GUS* seeds were transferred to the liquid 1× MS medium (4.44 g/L Murashige & Skoog modified basal medium powder with Gamborg vitamins, 0.5 g/L MES, 5 g/L sucrose, pH 5.7) in 96-well plates or 24-well plates with two seeds in 80 µL medium in each well or six seeds in 500 µL medium in each well for the GUS assay. The sterilized *A. thaliana pWER*::*FLS2-GFP* seeds were transferred to the square plates containing solid 0.5× MS medium (2.22 g/L Murashige & Skoog medium powder, 0.5 g/L MES, 5 g/L sucrose, 10 g/L agar, pH 5.7) with 3 rows of 20 seeds each row per square plate for the RGI assay. The 96-well plates or 24-well plates and the square plates were incubated in a short-day growth chamber for 9 days under a 10-h light/14-h dark cycle, with temperatures of 21°C during the day and 18°C at night, 70% relative humidity, and an LED white-light intensity of 250 µmol m⁻² s⁻¹. After germination, the seedlings were ready for use in the following steps in both assays.

### Preparation of bacteria

*Brevundimonas* sp. MF374 (NCBI RefSeq accession: GCF_900102995.1; JGI IMG accession: 2596583649) as a strong plant immunity suppressor was used as the target strain to investigate the mechanism of plant pattern-triggered immunity suppression in this study. This strain was isolated from Brassicaceae roots (mainly *Arabidopsis*) (*41*) and provided by Jeffery L. Dangl (University of North Carolina at Chapel Hill). For the Random Barcoded Transposon (RB-Tn) mutant library construction, *Escherichia coli* One Shot^TM^ Pir1 was purchased from Invitrogen. *E. coli* TransforMax EC100D pir+ was purchased from Lucigen. *E. coli* 5-alpha (DH5α) competent cells were purchased from New England Biolabs (NEB). Conjugation donor strain *E. coli* WM3064, originally constructed by William Metcalf (University of Illinois), was utilized for conjugation. All strains used in this study are listed in **Table S3**.

For the bacteria culturing, the freezer stocks of MF374 were streaked on the Luria-Bertani broth (Lennox) plates (used for MF374 WT and gene knockouts) and the LB plates supplemented with 50 μg/mL kanamycin (used for MF374 RB-Tn mutants, MF374 gene complements) and incubated at 28 °C for 48 hours. After incubation, a single colony of each strain was picked from a plate and inoculated into 5 mL of liquid LB medium and incubated at 28 °C for 24 hours under agitation at 250 rpm. After incubation, bacterial cultures were washed twice with 10 mM MgCl_2_ by centrifugation (5,000 × g, room temperature) and resuspension. Each bacterial pellet was resuspended in 1 mL of 10 mM MgCl_2_. The OD_600_ value of each strain was measured using NanoDrop One^c^ (Thermos Scientific). Cultures were diluted to required OD_600_ values using 1× MS medium.

### RB-Tn vector library construction

#### Construction of the nonbarcoded mariner vector pTGG31

Construction of the nonbarcoded mariner vector was adapted from a previous study (*22*). All plasmids used for RB-Tn vector library construction are listed in **Table S4,** and corresponding oligonucleotides are listed in **Table S5**. The nonbarcoded pTGG31 vector was assembled first. The ampicillin resistance gene was PCR amplified from pBAD-mTagBFP2 (Addgene), deposited by Vladislav Verkhusha (Albert Einstein College of Medicine), with primers DZ008 and DZ009. The R6K replication origin was PCR amplified from pW40, which was kindly provided by Yasuo Yoshikuni (DOE-JGI), with primers DZ010 and DZ011. PCR for the ampicillin gene was conducted using a 25 µL PCR system consisting of 12.5 µL Q5 High-Fidelity Master Mix (NEB), 0.5 µM primers, and 1 µL DNA under the following thermocycling conditions: 98 °C for 30 seconds, 30 cycles of 98 °C for 10 seconds, 62 °C for 30 seconds, and 72 °C for 30 seconds, and final extension at 72 °C for 2 minutes. PCR for the R6K origin was conducted similarly, except that 63 °C instead of 62 °C was used for annealing temperature in the 30 cycles. Two synthetic DNA constructs DZ023 (Twist Bio) and DZ024 (Genewiz) were synthesized including ends compatible with Golden Gate assembly.

A Golden Gate assembly reaction with DZ023 (Twist Bio), DZ024 (Genewiz), the ampicillin resistance gene, and the R6K origin were performed with BbsI (NEB) and NEBridge Ligase Master Mix (NEB) according to manufacturer instructions. The Golden Gate assembly reaction was purified with the DNA Clean & Concentrator kit (Zymo Research). The vector was transformed into the *E. coli* One Shot Pir1 (Invitrogen), then selected on LB plates with 100 μg/mL ampicillin. Vectors were miniprepped and Sanger sequenced with primers DZ001-DZ007 to check that the vector was assembled correctly.

#### Construction of the randomly barcoded mariner vector pTGG31 library

Construction of the barcoded mariner vector pTGG31 was adapted from a previous study (*22*). Briefly, the barcode containing oligonucleotide ofeba282, was PCR amplified with the primers ofeba285 and ofeba286 with Phusion polymerase (NEB), then purified with the DNA Clean & Concentrator kit (Zymo Research). The barcode PCR thermocycling conditions were changed to the following: 98 °C for 1 minute, 6 cycles of 98 °C for 10 seconds, 58 °C for 30 seconds, and 72 °C for 15 seconds, and a final extension at 72 °C for 5 minutes.

To construct the barcoded vector pTGG31 library, four Golden Gate assembly reactions were set up each with 1000 ng of the nonbarcoded pTGG31 and 40 ng of barcode PCR product with BsmBI (NEB) and NEBridge Ligase Master Mix (NEB) under the following thermocycling conditions: 30 cycles of 42 °C for 1 minute and 16 °C for 1 minute. The Golden Gate assembly reaction was purified with the DNA Clean & Concentrator kit (Zymo Research), then digested with BsmBI (NEB) under the following conditions: 55 °C for 2 hours and 80 °C for 20 minutes to remove any nonbarcoded vectors. The DNA was purified again with the DNA Clean & Concentrator kit (Zymo Research), transformed into the *E. coli* TransforMax EC100D pir+ (Lucigen), selected on LB plates with 100 μg/mL ampicillin. Vectors were miniprepped and Sanger sequenced with the primer DZ004 to check that the vector was assembled correctly.

#### Construction of the nonbarcoded Tn5 vector pDZ003

The nonbarcoded vector pDZ003 was derived from the nonbarcoded vector pTGG31. A PCR reaction was performed to amplify the portion of the nonbarcoded vector pTGG31 without the mariner specific promoter and transposase with primers DZ016 and DZ017 using 25 µL PCR system as described above under the following conditions: 98 °C for 30 seconds, 30 cycles of 98 °C for 10 seconds, 68 °C for 30 seconds, and 72 °C for 1 minute and 35 seconds, and a final extension of 72 °C for 2 minutes. The PCR product was treated with DpnI (NEB) for 1hr at 37 °C and for 20 minutes at 80 °C, then purified using the DNA Clean & Concentrator kit (Zymo Research).

A Gibson assembly reaction using the HiFi DNA Assembly Master mix (NEB) was performed with the PCR product and the synthetic DNA construct DZ025 (Twist Bio) and incubated at 50 °C for 1 hour. The Gibson assembly reaction was transformed into NEB DH5α Competent *E. coli* (NEB) and selected on LB plates with 100 μg/mL ampicillin. Site-directed mutagenesis with the Q5 site-directed mutagenesis kit (NEB) and primers DZ018 and DZ019 to replace the mariner specific IR near the barcode with the Tn5 specific IR using 25 µL PCR system as described above under the following conditions: 98 °C for 30 seconds, 25 cycles of 98 °C for 10 seconds, 72 °C for 30 seconds, and 72 °C for 2 minutes and 20 seconds, and a final extension of 72 °C for 2 minutes. The PCR product was treated with the KLD Master Mix (NEB), then transformed into the *E. coli* One Shot Pir1 (Invitrogen) and selected on LB plates with 100 μg/mL ampicillin. Vectors were miniprepped and Sanger sequenced with primers DZ004-007 and DZ020-022 to check that the vector was assembled correctly.

#### Construction of the randomly barcoded Tn5 vector pDZ003 library

Construction of the vector pDZ003 library was completed as described with the construction of the barcoded vector pTGG31 library, but the Golden Gate assembly reaction was set up with the nonbarcoded pDZ003 instead of the nonbarcoded pTGG31.

### RB-Tn mutant library construction

#### WM3064 Library Construction

The vector pTGG31 library and vector pDZ003 library were transformed into diaminopimelic acid (DAP)-auxotrophic *E. coli* conjugation strain WM3064 with electroporation. A mixture of 10 µL of a vector library and 90 µL of WM3064 cells were electroporated in the GenePulser Xcell (BioRad) in 1 mm cuvettes with the following conditions: 1.8 kV, 25 μF, 200 Ω. Transformed WM3064 were selected on LB agar plates containing 100 μg/mL ampicillin, 50 μg/mL kanamycin, 50 μg/mL erythromycin, and 300 μM DAP. Colonies were scraped into LB media containing 100 μg/mL ampicillin, 50 μg/mL kanamycin, 50 μg/mL erythromycin, and 300 μM DAP. The 1.5 mL freezer stocks were made containing a final concentration of 20% glycerol and frozen at -80 °C.

#### MF374 RB-Tn mutant Library Construction

Fresh culture of *Brevundimonas* sp. MF374 was prepared as addressed above. The WM3064_pTGG31 and WM3064_pDZ003 freezer stocks were each added into 50 mL of LB medium containing 100 μg/mL ampicillin, 50 μg/mL kanamycin, 50 μg/mL erythromycin, and 300 μM DAP and incubated at 37 °C for 1.5 hours. For the MF374 mutant library construction, MF374 and both the WM3064 libraries were washed twice with LB media, all resuspended in LB media, then combined in a volumetric ratio of 2:1 MF374 to the WM3064 library and washed again. Combined bacteria resuspended in 100 μL LB and were plated on LB agar plates containing 300 μM DAP and incubated at 28 °C overnight. Conjugations were scraped into 2 mL LB media, diluted 1:5 in LB media, then plated on LB agar plates containing 50 μg/mL kanamycin and incubated at 28 °C for two days. Colonies were scraped into LB media containing 50 μg/mL kanamycin and split into 1.5 mL freezer stocks containing a final concentration of 20% glycerol. A total of over 100,000 colonies of each library were collected.

#### RB-TnSeq Library Construction

MF374 mutant libraries were characterized using RB-TnSeq as previously described (*42*). In short, genomic DNA was fragmented with a Covaris LE220 focused-ultrasonicator. The final fragmented and size-selected genomic DNA (400-500 bp) was quantified with a Qubit dsDNA HS assay kit (Invitrogen). Nspacer_barseq_universal was used for the transposon-specific primer. All oligonucleotides used for RB-TnSeq library preparation are listed in **Table S4**. The final PCR product was quantified with a Qubit dsDNA HS assay kit (Invitrogen). RB-TnSeq samples were run on a 2% agarose gel with the GeneRuler 1kb Plus DNA Ladder (Thermo Scientific) and stained with SYBR Safe DNA Gel Stain (Invitrogen). RB-TnSeq samples were also checked for the average size (450 bp) and concentration on the Agilent Bioanalyzer 2100 (Agilent). The NEBNext Ultra II DNA Library Prep Kit for Illumina (NEB) was used to construct two pTGG31 and two pDZ003 RB-TnSeq libraries following the kit instruction. The four RB-TnSeq libraries were pooled using equimolar DNA from each library. The constructed RB-TnSeq library was sequenced with NovaSeq 6000 (Illumina) with PE150 service at Genewiz (Azenta Life Sciences).

#### RB-TnSeq Data Analysis

RB-TnSeq analysis followed the TnSeq analysis pipelines (github.com/m-jahn/TnSeq-pipe), which is based on RB-Tnseq script in Feba repository (bitbucket.org/berkeleylab/feba/src/master/) (*42*, *43*). Briefly, the Tn insertion sites across the whole MF374 genome were identified. Through genome mapping, the genes with inserted Tns and unique barcodes were annotated based on the MF374 genome annotation (accession: GCF_900102995.1) from the NCBI RefSeq. The unique mapped barcodes for all the insertion sites were summarized. The library used for the following mutant screening was selected based on comparison between pTGG31 and pDZ003 RB-TnSeq library in the number of the mapped unique barcodes.

#### RB-Tn mutant screening preparation

*A. thaliana pCYP71A12*::*GUS* seedlings in 96-well plates were prepared as described above (*19*). The MF374_pDZ003 RB-Tn mutant library was plated on LB agar plates with 50 μg/mL kanamycin and grown at 28 °C for 2-4 days. Individual colonies were picked to be placed into each well of the 96-deep well plates with 1 mL LB medium with 50 μg/mL kanamycin and grown at 28 °C for 24 hours. Bacteria were washed twice with 10 mM MgCl_2_ by centrifugation at 5,000 × g, then resuspended in 500 μL MS medium. One microliter of the mutants from the 96-deep well plates were added into wells of the 96-well plates containing the *A. thaliana pCYP71A12*::*GUS* seedlings and placed in the short-day growth chamber for 14 hours. The 96-deep well plates containing the mutant bacteria were stored with a final concentration of 20% glycerol in each well and frozen at -80 °C.

#### GUS assay for mutant screening

The next day, 1 μM of flg22-*Pa* (from *Pseudomonas aeruginosa*) peptide was added into each well and the plates were placed back into the same growth chamber for 5 hours. Seedlings were washed once with 50 mM sodium phosphate buffer at pH 7, then 80 μL of freshly prepared GUS substrate solution was added to each well and incubated at 37 °C in the dark for 16 hours. GUS substrate was made with 50 mM sodium phosphate buffer at pH 7, 10 mM EDTA, 0.5 mM K_4_[Fe(CN)_6_], 0.5 mM K_3_[Fe(CN)_6_], 0.5 mM X-Gluc, and 0.01% Silwet L-77. GUS substrate solution was replaced with 80 μL of a 3:1 ethanol: acetic acid solution. Plates were left at 4 °C overnight, then the ethanol and acetic acid solution was replaced with 80 μL of 95% ethanol for imaging. Roots were imaged on an AmScope LED-114S with the camera AmScope FMA050 (Amscope). Roots were manually observed for the tips with blue stain. A white root tip with MF374 WT was used as negative control due to the flg22-*Pa*-triggered plant immunity suppressed by this strain. A root tip with blue stain with no bacteria (NB) was used as positive control due to plant immune system activated by flg22-*Pa*. The root tip with the blue stain suggests the mutant strain loses the ability to suppress plant immunity. The RB-Tn mutant screening was conducted on 4,032 mutant strains.

#### Arbitrary PCR and Sanger Sequencing

After mutant screens, five RB-Tn mutant candidates were obtained. The mutant candidates were subjected to two rounds of arbitrary PCR following a protocol from O’Toole et al. (*44*) and Teixeira et al. (*19*) with some modifications. This first PCR was conducted using a forward primer (JMC525) targeting on kanamycin marker from the inserted Tn and a reverse primer (ARB6) (*44*) arbitrarily targeting sequence on the mutant genome under a touchdown PCR program: 98 °C for 3 min, 15 cycles of 98 °C for 30 seconds, 50 °C (1 °C decreased per cycle) for 45 seconds, and 72 °C for 3 min, and another 20 cycles of 98 °C for 30 seconds, 35 °C for 45 seconds, and 72 °C for 3 min, and a final extension of 72 °C for 5 minutes. Afterward, the second round of PCR was performed on the PCR product using a forward primer (JMC567) nested inside the kanamycin resistance gene and a reverse primer (ARB2) (*44*) randomly binding on a site in the MF374 mutant genome under the same touchdown PCR program but with the initial annealing temperature set to 81 °C. Genomic DNA of the mutants was extracted using the DNeasy Blood & Tissue Kit (Qiagen). The touchdown PCR was employed to increase the chance of the arbitrary primers binding on the genome (*45*). PCR products were submitted to Genewiz (Azenta Life Sciences) for Sanger sequencing after being purified using DNA Clean & Concentrator kit (Zymo Research). After sequencing, the sequence was blasted against pDZ003 sequence to identify the part of the sequence with the end of IR, then the rest of the sequence was blasted against MF374 genomes with accessions 2596583649 and GCF_900102995.1 in JGI IMG and NCBI RefSeq databases, respectively, to identify the genes inserted by Tns.

#### Normalized GUS assay for RB-Tn mutant candidate validation

All the five RB-Tn mutant candidates that lost the flg22-*Pa*-triggered immune attenuation were validated using a normalized GUS assay. The RB-Tn mutant candidate cultures were prepared as described above. The cultures were diluted using 1× MS medium from 10⁻² to 10⁻⁴ of OD_600_. Then, 500 µL of each dilute was transferred to a well in 24-well plates with 9-day-old *A. thaliana pCYP71A12*::*GUS* seedlings (2-6 seedlings after germination). The plates were incubated in the short-day growth chamber for 14 hours. After incubation, flg22-*Pa* were added into two wells of each plant-mutant co-culture for 1 and 0.1 µM, respectively. The plates were placed back in the same growth chamber for 5 hours. Afterward, seedlings were submitted to the GUS substrate for staining and washing process, and roots were imaged following the same protocol described above except for 500 µL of the GUS substrate, 3:1 ethanol to acetic acid solution, and 95% ethanol used in each well of 24-well plates instead of 80 µL.

#### RGI assay for RB-Tn mutant candidate validation

The prepared RB-Tn mutant cultures at OD_600_ = 0.01 were spread on 0.5× MS square plates without 0.5% sucrose added. The plates contained 1 μM flg22-*Pa* group and mock (sterile DI-water) group. Eight 9-day-old seedlings of *A. thaliana pWER*::*FLS2-GFP* were transferred to bacteria-spread 0.5× MS square plates. The end of root tips was labeled with short straight lines. The plates were incubated in a short-day growth chamber for 9 days under a 10-h light/14-h dark cycle, with temperatures of 21°C during the day and 18°C at night, 70% relative humidity, and an LED white-light intensity of 250 µmol m⁻² s⁻¹. After incubation, the plates were photographed, then elongation of a root in each photo was measured starting from the labeled line to the new end of the root tip using ImageJ. The differences in root elongation between WT and RB-Tn mutant candidates were evaluated to assess the loss of attenuation of flg22-*Pa*-triggered RGI by the mutants.

#### Gene knockout vector construction

To validate gene function of the immune attenuation phenotypes for the identified *ppiD* and *tonB*, two single gene knockout mutants, Δ*ppiD* and Δ*tonB1::GmR*, and a double gene mutant (Δ*tonB1::GmR* Δ*ppiD*) were constructed. As *ppiD* has four-nucleotide (ATGA) overlaps with the downstream gene cluster *trpEXGDC*, a single gene knockout (Δ*trpEXGDC*) and two double gene knockouts (Δ*ppiD*-*trpEXGDC*) and (Δ*tonB1::GmR* Δ*trpEXGDC*) were constructed as shown in **Table 1**. The gene knockout vectors were designed to contain the pMO130 backbone (*17*), 5’ flanking region and 3’ flanking region of the target gene for knockout. For each vector, the two 1000 bp flanking regions were amplified from MF374 DNA using two sets of primer pairs (**Table S8**), which were designed using the NCBI primer design tool and synthesized in IDT. The pMO130 backbone was obtained through amplifying the vector with primers JMC203 and JMC204 using 25 µL PCR system as described above under the following conditions: 98 °C for 30 seconds, 32 cycles of 98 °C for 10 seconds, 60 °C for 10 seconds, and 72 °C for 2 minute, and a final extension of 72 °C for 5 minutes (*49*). Gibson assembly was constructed for assembling the backbone, 5’ flanking region, and 3’ flanking region using HiFi DNA Assembly Master mix (NEB) at 50 °C for 1 hour. All gene knockout vectors were listed in **Table S7**.

#### *E. coli* DH5*α* Transformation for gene knockouts

Each assembled vector was diluted 1:3 with sterile nuclease-free water. Five microliters of the vector dilute was mixed with 50 µL *E. coli* DH5α competent cells. The mixture was placed on ice for 30 min, moved to 42 °C water batch for heat shock for 30 seconds, and put on ice for 5 min. The processed cells were incubated in 1 mL Super Optimal Broth (SOB) at 37 °C for 3 hours. The culture was spread on a LB plate, and the plate was incubated at 37 °C for 24 hours. Six colonies were picked from the plate, and each colony was inoculated in a culturing tube with 5 mL of LB broth. The inoculated broth was incubated at 37 °C for 24 hours. After incubation, each vector was extracted from the cultures using a ZymoPURE Plasmid Miniprep Kit (Zymo Research). The extracted plasmids were submitted to Genewiz (Azenta Life Sciences) for the whole plasmid sequencing service. The sequenced plasmids were aligned with the designed vector sequence using the SnapGene software (version 8.2).

#### *E. coli* WM3064 Transformation for gene knockouts

The sequence confirmed vectors were transformed into the DAP-auxotrophic *E. coli* WM3064 competent cells using the same processing as used for vectors transformed into *E. coli* DH5α described above. After the transformed WM3064 grew in SOB at 37 °C for 3 hours, the culture was spread onto the LB agar plate with 300 µM DAP. The plates were incubated at 37 °C for 24 hours. After incubation, six colonies from one plate were picked for checking the success of vector transformation by conducting colony PCR, which used 25 µL reaction system as described above under the following conditions: 98 °C for 10 min, 30 cycles of 98 °C for 30 seconds, annealing temperature (AT) (**Table S8**) for 45 seconds, and 72 °C for 30 seconds, and a final extension of 72 °C for 5 minutes with a pair of primers (**Table S8**) targeting the 5’ flanking region and kanamycin resistant gene.

#### Biparental mating for gene knockouts

Once the vector was confirmed in the cell, the confirmed WM3064 was cultured in liquid LB medium with 300 µM DAP at 37 °C for 24 hours, and MF374 was cultured in liquid LB medium at 28 °C for 24 hours. Then, the two cultures were washed two times with 10 mM MgCl_2_ by centrifugation at 5,000 g for 10 min at room temperature. Fifty microliters of each culture were mixed well in a 1.5 mL centrifuge tube. A 50 µL of the mixture was added onto a LB agar plate with 0.5 mM DAP, which was incubated at 28 °C for 24 hours. Then, the puddle was scraped after 1 mL of LB liquid medium was added. The scraped cell culture was diluted 10-fold. One hundred microliters of the origin and dilute were spread on LB agar plates and incubated at 28 °C for 48 hours. After incubation, six colonies were picked for colony PCR to check the success of homogeneous recombination. The PCR program was the same as described above with primers (**Table S8**) targeting a site outside of the 5’ flanking region and the kanamycin resistant gene.

#### Sucrose counter selection for gene knockouts

The colony with successful homogeneous recombination was grown in 5 mL of liquid LB media with 0.5 mM IPTG and 10% sucrose at 28 °C for 48 hours. The culture was streaked on an LB plate with 0.5 mM IPTG and 10% sucrose and incubated at 28 °C for 48 hours. Afterward, 95 colonies were picked with sterile toothpicks and transferred into a 96-well plate with 100 µL sterile water in each well. Then, 10 µL of each colony solution was transferred to one well of another 96-well plate with 90 µL of LB liquid media with kanamycin at a final concentration of 50 µg/mL, which was incubated at 28 °C for 24 hours. A negative control of 10 µL of sterile DI-water was added in the last well. After incubation, wells with the media showing on turbidity indicate the gene knockout candidates. Then, colony PCR was conducted using gene knockout candidates with primers (**Table S8**) targeting a site outside of the 5’ flanking region and a site outside of the 3’ flanking region. The colony PCR program was the same as described above except for different ATs used (**Table S8**). The confirmed gene knockouts were grown in liquid LB medium at 28 °C for 24 hours. The incubated culture was washed twice with 10 mM MgCl_2_ and resuspended with 1 mL of liquid LB medium. The culture was transferred into a 2 mL cryogenic tube to make a culture stock containing a final concentration of 20% glycerol and stored at -80 °C.

#### Gene complement vector construction

To further validate the gene function, six gene complements were constructed using gene knockout cells and complement vectors, including Δ*ppiD* / pBBR1::*ppiD*, Δ*tonB1*::*Gm^R^* / pBBR1::*tonB1*, Δ*trpEXGDC* / pBBR1::*trpEXGDC*, Δ*ppiD*-*trpEXGDC* / pBBR1::*ppiD*, Δ*ppiD*-*trpEXGDC* / pBBR1::*trpEXGDC*, Δ*ppiD*-*trpEXGDC* / pBBR1::*ppiD*-*trpEXGDC* as shown in **Table 1**. Each of the complement vectors contains a pBBR1 backbone and the complemented gene. Primer pairs for amplification of the complemented genes were designed based on the MF374 genome and synthesized in IDT. Gibson assembly was conducted to assemble the backbone and the complemented gene in a PCR machine at 50 °C for 1 hour. The PCR system contained assembly master mix, the pBBR1 backbone, and the complement gene. All gene complement vectors were listed in **Table S7**.

#### *E. coli* DH5*α* and WM3064 transformation for gene complements

By following the same protocol of the gene knockout except for the sucrose selection step described above, a complement vector was transformed into competent *E. coli* DH5α cells. After enrichment, the vector was extracted using ZymoPURE Plasmid Miniprep Kit (Zymo Research) and submitted to Genewiz (Azenta Life Technologies) for the whole plasmid sequencing service. Then, the confirmed complement vector was transformed into DAP-auxotrophic *E. coli* WM3064 competent cells.

#### Conjugation for gene complements

Through conjugation, the complement vector was passed from the transformed WM3064 into the gene knockout. Colony PCR following the same program as described above except for ATs (**Table S8**) was carried out to test the success of gene knockout transformation with primers (**Table S8**) targeting a site on the complement gene and a site on the kanamycin resistance gene. The confirmed cell was grown in LB liquid media with 50 µg/mL kanamycin at 28 °C for 24 hours. The culture was washed and stored in 2 mL of LB liquid media with 50 µg/mL kanamycin and 20% glycerol at -80 °C.

#### Normalized GUS and RGI assays for gene function validation

Normalized GUS assay follows the same procedure as described above, except for bacteria used from 1 to 10^-5^ of OD_600_ instead of 10^-2^ to 10^-4^ and immune elicitors used at 1 µM instead of 0.1 and 1 µM. The RGI assay follows the same procedure as described above.

#### Root colonization assay

The root colonization assay was based on the normalized RGI assay. After the seedlings were incubated for 9 days in the short-day growth chamber, the roots were cut from the plants on each plate and placed into a 1.5 mL tube. The root sample was washed with 1 mL of 10 mM MgCl_2_ twice through a vortex (Fisherbrand) at 3,000 rpm for 15 seconds each time. The root sample was dried on a sterilized filter paper. After drying, the root sample was transferred to a 2 mL sterile screw tube with 3 glass beads (4 mm). Each tube was weighed before and after the root sample added, and then the root sample weight was calculated. Four hundred microliters of 10 mM MgCl_2_ were added into the tube. The tubes were placed on a bead beater (MP) for root tissue homogenization at 6 m/sec for 40 seconds. Then, 600 µL of 10 mM MgCl_2_ were added into the tube. The root sample was serially diluted 10-fold. The dilutions were plated on LB agar plates and incubated at 28 °C for 48 hours. After incubation, the colonies were enumerated using a gel imager (SYNGENE) with software GeneSys for imaging and software GeneTools for colony counting.

#### Bacterial growth curve measurement

MF374 wild type strain, three single gene deletion strains (Δ*ppiD*, Δ*tonB1*::*GmR*, and Δ*trpEXGDC*), three complemented strains (Δ*ppiD* / pBBR1::*ppiD*, Δ*tonB1*::*Gm^R^* / pBBR1::*tonB1*, and Δ*trpEXGDC* / pBBR1::*trpEXGDC*), and five Tn-mutated strains (*ppiD*::Tn, *tonB1*::Tn, *RS09010*::Tn, *purD*::Tn, and *pmbA*::Tn) were prepared as described above. Each washed strain was prepared at OD_600_ = 0.01 in 1× MS medium containing 1% Casamino acids without and with 1 mM Tryptophan. Two hundred microliters of each culture were transferred into a well of a 96-well plate. Growth curve of each strain in the plate was monitored by BioTek Synergy H1 plate reader (Agilent) with condition setting as OD_600_ value of each strain at 28 °C measured every two hours for 72 hours. The experiment was performed in three biological replicates with three technique replicates.

#### Designed GUS assay panel

A GUS assay panel was designed to evaluate the contribution of different parts in the plant-bacteria co-culture system to the immune attenuation phenotype of MF374, resulting in five distinct GUS assays (referred to here as A1-A5). The differences among the five assays appear after 14-hour incubation for MF374 WT and gene knockouts growing in 1× MS. The first assay (A1) is the regular GUS assay as shown above, which contained plant-bacteria co-culture system. The second one (A2) used the co-culture medium for the following GUS steps. The third one (A3) transferred the bacteria-colonized seedlings from a well washed with 10 mM MgCl_2_ twice (washed seedings) into a well in a new 24-well plate, where fresh 1x MS medium with 1 µM flg22-*Pa* was added, followed by incubation in the short-day growth chamber for 5 hours. The fourth one (A4) applied a 0.22 µm syringe filter to filter the 500 µL co-culture medium into a new well to get the cell-fresh supernatant, and new sterile seedlings were added, followed by flg22-*Pa* addition and incubation for 5 hours in the short-day growth chamber. The fifth one (A5) transferred the co-culture medium to a new well, where flg22-*Pa* was added at 1 µM. After 5-hour incubation in the short-day growth chamber, the cell-flg22-*Pa* pre-interacting medium was filtered using a 0.22 µm syringe filter into a new well with sterile seedlings in it. The following step was the same for the five assays, which was to use the prepared GUS substrate for GUS staining and washing and root imaging using an AmScope LED-114S with the camera AmScope FMA050 (Amscope) as described above.

#### Immune suppression assays using additional elicitors

First, we tested if five different immune elicitors, including 1 µM flg22-*Pa* (as a control) flg22-*Dja* (from *D. japonica* MF79), AtPEP1, and elf18 and 200 µg/mL chitin could activate plant immune in *A. thaliana pCYP71A12*::*GUS* plants using the GUS assay. Following this, we evaluated if 0.1 µM AtPEP1, and elf18 and 20 µg/mL chitin could activate immune system in *A. thaliana* Col-0 plants using the RGI assay, and if the elicitors-triggered immunity could be suppressed by MF374 WT from 0.01 to 1 of OD_600_. Moreover, 1 µM flg22-*Dja* was used to activate the immune system in *A. thaliana pCYP71A12*::*GUS* plants, and the immunity suppression by MF374 WT and the loss of the immunity suppression by gene knockouts Δ*ppiD*, Δ*tonB1*::*GmR*, Δ*tonB1*::*GmR* Δ*ppiD*, and Δ*trpEXGDC*, at OD_600_ = 0.01 were assessed. For the other three elicitors, 0.1 µM AtPEP1, elf18, and 20 µg/mL chitin were applied to activate the immune system in *A. thaliana* Col-0, and the loss of immunity suppression by gene knockouts Δ*ppiD*, Δ*tonB1*::*GmR*, Δ*tonB1*::*GmR* Δ*ppiD*, and Δ*trpEXGDC*, at OD_600_ = 1 and 0.1 only used for AtPEP1 group were assessed. The detailed RGI procedure was described above. The peptide sequences of all immune elicitors are listed in **Table S9**.

#### Construction of phylogenetic trees for TonB and TonB-dependent transporter (TBDT) proteins

Five Annotated genome assemblies, including MF374 (GCF_900102995.1), MF79 (GCF_000620485.1), WCS417 (GCF_000698265.1), *Rhodanobacter* sp. R179 (GCF_001428465.1), and *E. coli* K12 (GCF_000005845.2), were searched in NCBI, respectively. Annotated genes were view. For each bacterial strain, “TonB” was searched through the annotated genes under the Filters option. FASTA files of all the displayed TonB-related proteins for each bacterial strain were downloaded. All the TonB-related protein sequences from five strains were split into TonB, including TonB energy transducer and TonB family proteins, and TBDT, including TonB-dependent transporter (annotated as receptor in RefSeq) related sequences groups. For each group, SnapGene was used to align multiple protein sequences using the MAFFT alignment method. The aligned protein file was opened in MEGA12 for phylogenetic analysis obtain the initial phylogenetic tree using an evolutionary analysis of maximum likelihood method with Jones-Taylor-Thornton (JTT) model. The final phylogenetic trees for TonB and TBDT were made, respectively, using the R packages ggtree, ggtreeExtra, and ggplot2.

#### Statistical data analysis

All experiments were conducted with at least three biological replicates. All statistical data analysis was performed using R software (version 4.5.1). Nonparametric Dunn’s test (*46*) with the Benjamini-Hochberg method for *P*-value adjustment was applied for multiple comparison analysis following a significant Kruskal-Wallis test using R package FSA (version 0.10.1). Difference with *P* < 0.05 is considered statistically significant. Detailed statistical data analysis for different experiment data was addressed in individual figure captions.

## Supplementary Materials

Figs. S1 to S9

Tables S1 to S11 (separate files)

## Acknowledgments

We sincerely thank J.L. Dangl for bacterial strains *B.* sp. MF374, W. Metcalf for bacterial strain *E. coli* WM3064, V. Verkhusha for vector pBAD-mTagBFP2, and Y. Yoshikuni for vector pW40.

## Author contributions

Conceptualization: J.M.C. Methodology: J.M.C. and C.L. Investigation: C.L., D.Z., A.Z., J.G. Visualization: C.L. Supervision: J.M.C. Writing— original draft: C.L. Writing—review & editing: J.M.C. and C.L. All authors reviewed and approved the final manuscript.

## Data and materials availability

All data and codes needed to evaluate and reproduce the results in the paper are present in the paper, the Supplementary Materials, and/or https://github.com/LZC0034/MF374_RB-TnSeq. The raw RB-TnSeq data was deposited to the NCBI database under accession number PRJNA1550221.

