## Supplementary Materials for "Genetic determinants of *Brevundimonas*-mediated attenuation of plant pattern-triggered immunity"


**This PDF file includes:**

Figs. S1 to S9

Tables S1 to S11 (separate files)

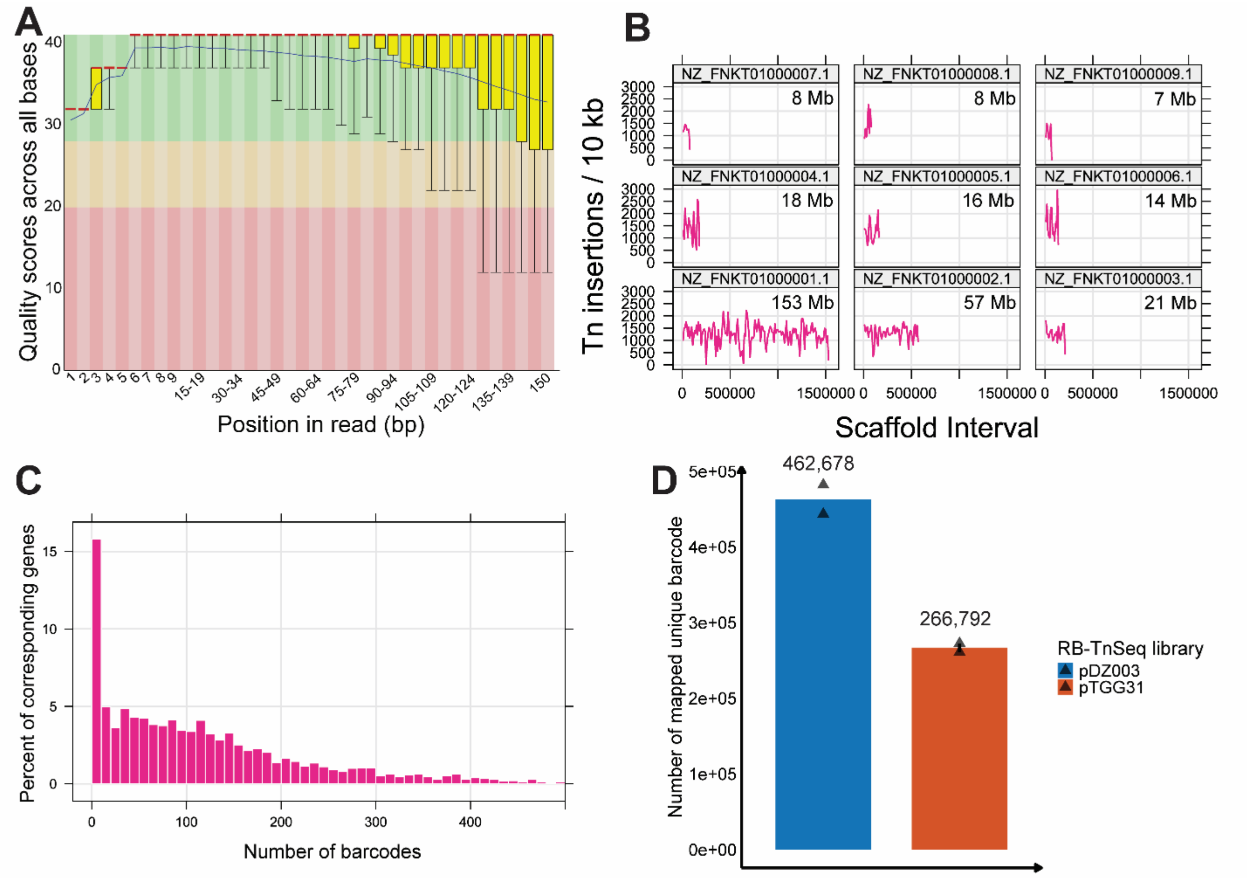


**Fig. S1. Analysis of the pDZ003 and pTGG31 RB-TnSeq mutant libraries in *Brevundimonas* sp. MF374. (A)** Per-base quality scores for the pDZ003 random barcoded transposon (RB-TnSeq) library in MF374. **(B)** Distribution of transposon insertions per 10 kb across the nine scaffolds of the *Brevundimonas* sp. MF374 genome in the pDZ003 library. The number of Tn insertions per 10 kb across the genome is 1293 ± 161. **(C)** Distribution of mapped barcodes per gene in the pDZ003 library. **(D)** Comparison of the average number of mapped unique barcodes between the pDZ003 and pTGG31 RB-TnSeq libraries. Two biological replicates of each library were sequenced and analyzed together.

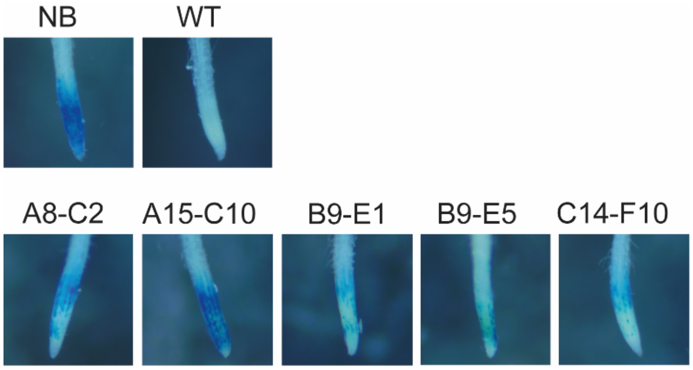


**Fig. S2. Identification of five candidate *Brevundimonas* sp. MF374 transposon mutants with impaired immune attenuation.** Root tip images of five candidates obtained from screening of 4,032 Tn-insertion mutants of *Brevundimonas* sp. MF374 using the *A. thaliana pCYP71A12*::*GUS* reporter line. Blue staining indicates flg22-Pa triggered immune activation. Candidate labels indicate the screening batch (first letter), 24-well plate number, and well position (final letter and number). The five candidates correspond to A8-C2 (*ppiD*::Tn), A15-C10 (*tonB1*::Tn), B9-E1 (*RS09010*::Tn), B9-E5 (*purD*::Tn), and C14-F10 (*pmbA*::Tn). NB: no bacteria; WT: MF374 wild type.

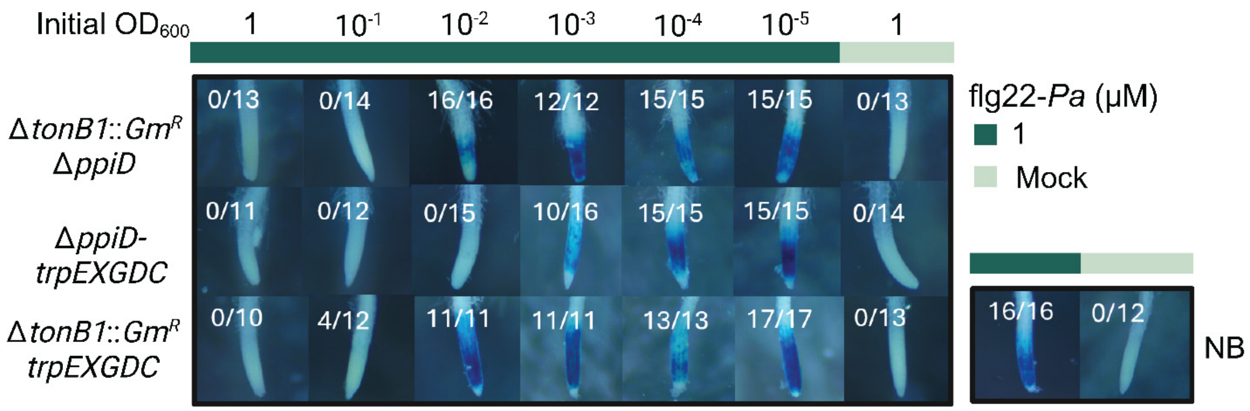


**Fig. S3. Analysis of double mutants supports a dominant role for *tonB1* in MF374-mediated immune attenuation.** GUS reporter assays of *Brevundimonas* sp. MF374 Δ*tonB1*::*Gm*$^R$ Δ*ppiD*, Δ*ppiD-trpEXGDC*, and Δ*tonB1*::*Gm*$^R$ Δ*trpEXGDC* mutants using *A. thaliana pCYP71A12*::*GUS* seedlings. Bacteria were applied at initial $OD_{600}$ values from 1 to $10^{-5}$ with 1 μM flg22-*Pa*. Fractions indicate the number of blue-stained root tips relative to the total number examined across three biological replicates. Corresponding single-mutant results are shown in Fig. 2A. The no-bacteria (NB) control images, with and without flg22-*Pa*, are also presented in Fig. 2A. NB: no bacteria; Mock: sterile deionized water control.

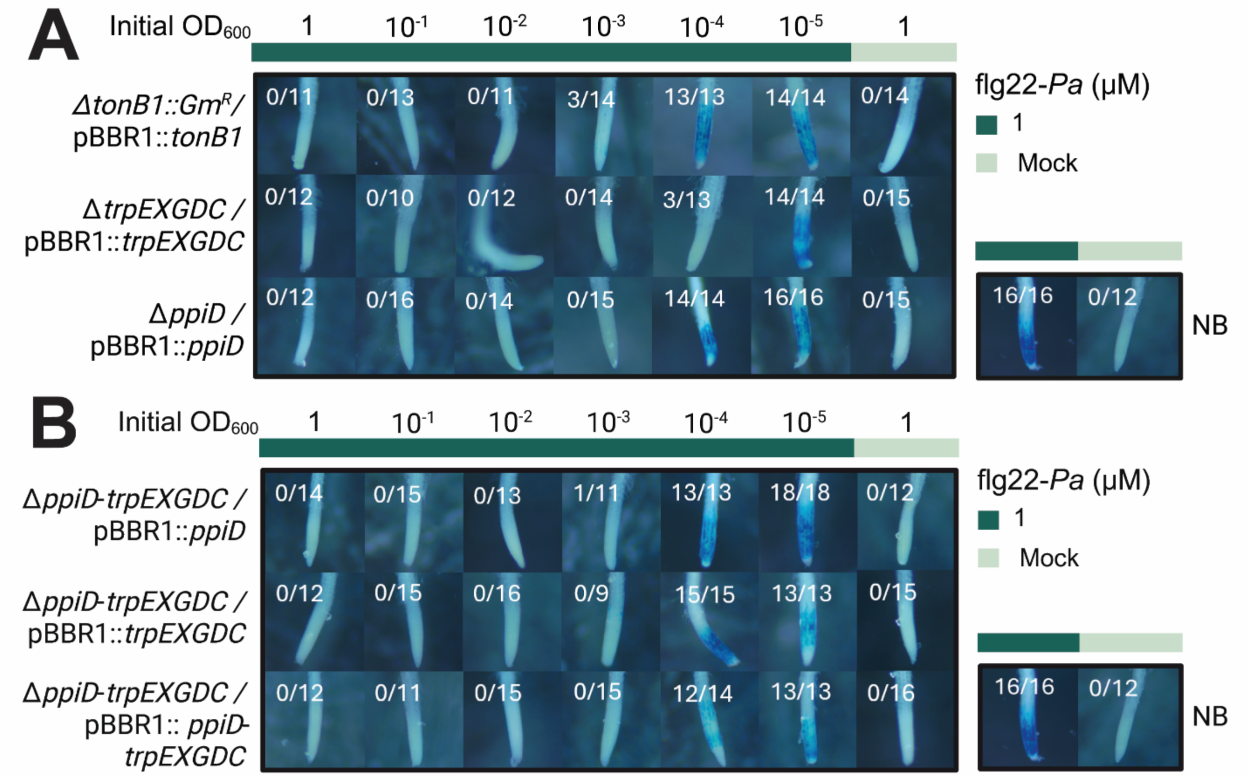


**Fig. S4. Gene complementation in *Brevundimonas* sp. MF374 mutants restores attenuation of flg22-*Pa*-triggered plant immunity to varying degrees. (A)** GUS reporter assays of complemented MF374 single mutants (Δ*ppiD* / pBBR1::*ppiD*, Δ*tonB1*::*Gm*$^{R}$ / pBBR1::*tonB1*, and Δ*trpEXGDC* / pBBR1::*trpEXGDC*). **(B)** GUS reporter assays of the Δ*ppiD-trpEXGDC* mutant complemented with *ppiD*, *trpEXGDC*, or *ppiD-trpEXGDC.* Assays used the *A. thaliana pCYP71A12*::*GUS* reporter line with bacteria at initial $OD_{600}$ of 1 to $10^{-5}$ and 1 µM flg22-*Pa*. Fractions indicate the number of blue-stained root tips relative to the total number examined across three biological replicates. The no bacteria (NB) control images, with and without 1 µM flg22-*Pa,* are also presented in Figure 2A. NB: no bacteria. Mock: sterile deionized water control.

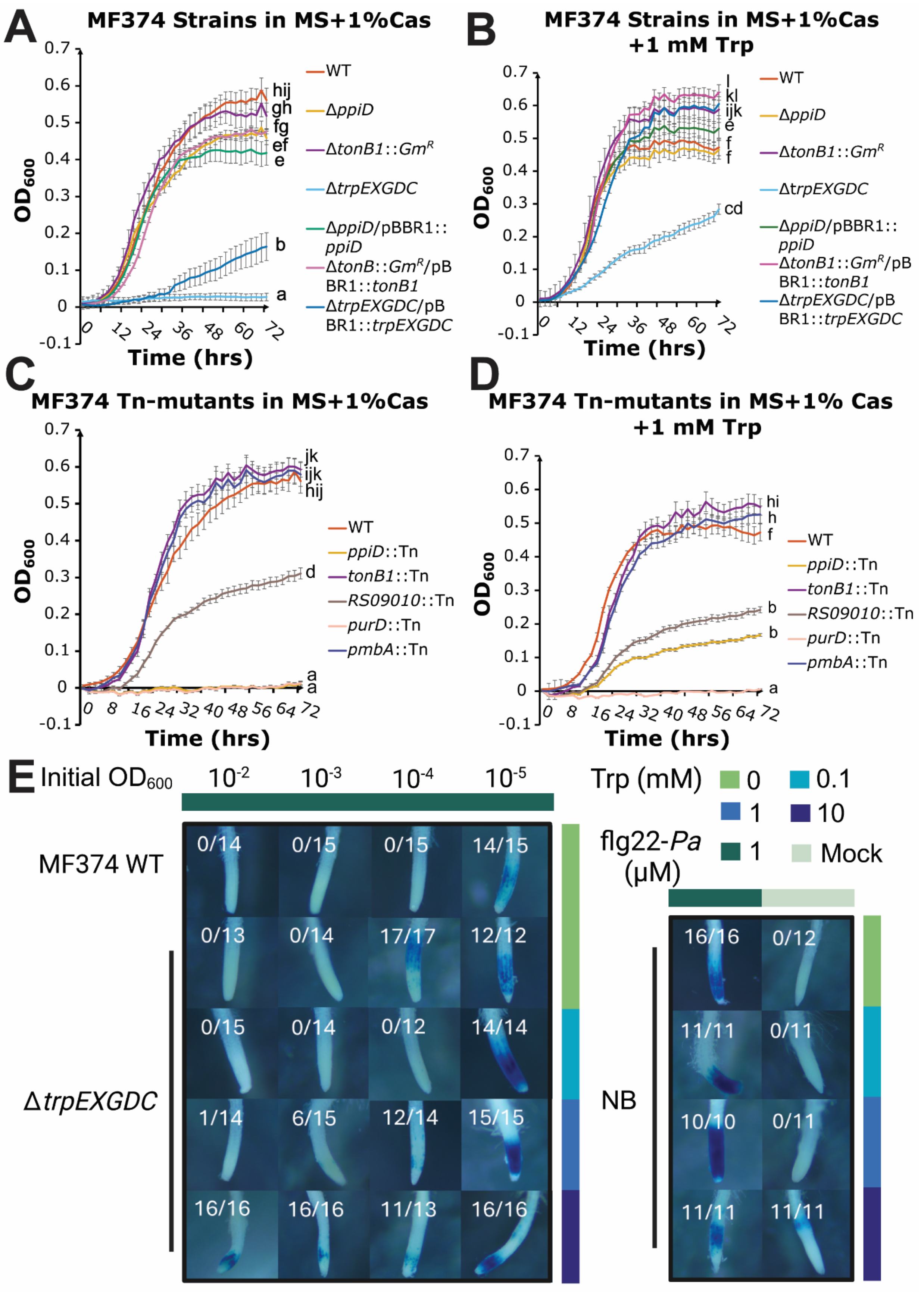
A
MF374 Strains in MS+1%Cas
B
MF374 Strains in MS+1%Cas +1 mM Trp
C
MF374 Tn-mutants in MS+1%Cas
D
MF374 Tn-mutants in MS+1% Cas +1 mM Trp
OD600
Time (hrs)
WT
ΔppiD
ΔtonB1::GmR
ΔtrpEXGDC
ΔppiD/pBBR1::ppiD
ΔtonB::GmR/pBBR1::tonB1
ΔtonB1::GmR/pBBR1::tonB1
ΔtrpEXGDC/pBBR1::trpEXGDC
ppiD::Tn
tonB1::Tn
RS09010::Tn
purD::Tn
pmbA::Tn
E
Initial OD600
10-2
10-3
10-4
10-5
Trp (mM)
0
0.1
1
10
flg22-Pa (μM)
1
Mock
MF374 WT
ΔtrpEXGDC
NB

**Figure S5. Disruption of tryptophan synthesis genes *trpEXGDC* impairs bacterial fitness of *Brevundimonas* sp. MF374. (A–D)** Growth curves of MF374 wild type (WT), gene deletion and complemented strains **(A,B)**, and transposon mutant strains **(C, D)** in 1× MS medium supplemented with 1% Casamino acids (Cas), without (**A, C**) or with (**B, D**) 1 mM tryptophan (Trp). Cultures were grown at 28°C for 72 h. $OD_{600}$ was measured every 2 h using a plate reader. Data are presented as mean ± SD from three biological replicates, each with three technical replicates. Different letters indicate significant differences in $OD_{600}$ at 72 h ($P < 0.05$; one-way ANOVA followed by Tukey's HSD test). **(E)** GUS reporter assays of MF374 WT and Δ*trpEXGDC* mutant using *A. thaliana pCYP71A12*::*GUS* seedlings with 0, 0.1, 1, or 10 mM tryptophan and 1 μM flg22-*Pa*. Bacteria were applied at initial $OD_{600}$ from $10^{-2}$ to $10^{-5}$. Fractions indicate the number of blue-stained root tips relative to the total number examined across three biological replicates. WT: wild type; NB: no bacteria; Mock: sterile deionized water.

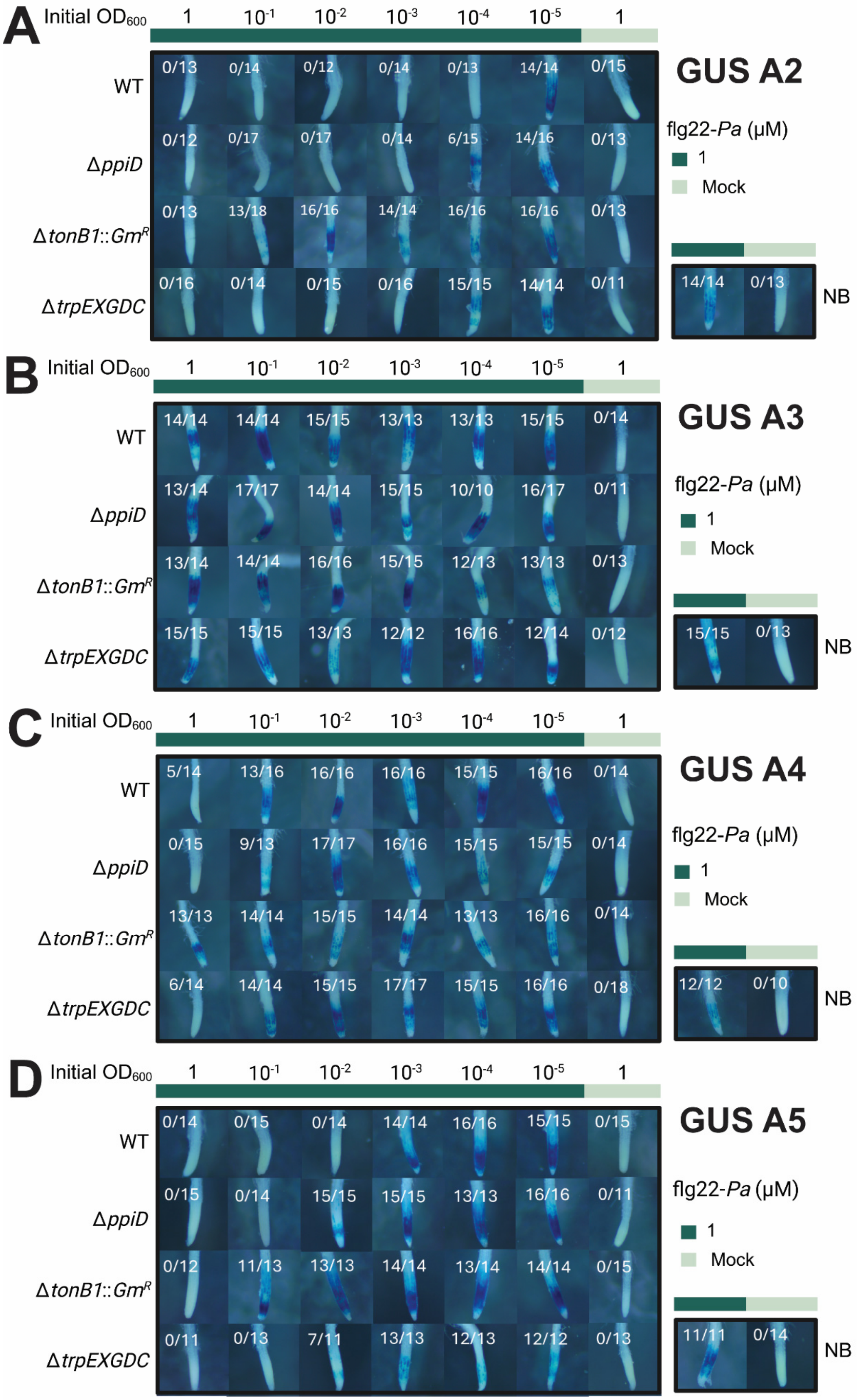
A Initial OD600 1 10-1 10-2 10-3 10-4 10-5 1
GUS A2
flg22-Pa (μM)
1
Mock
NB
WT
ΔppiD
ΔtonB1::GmR
ΔtrpEXGDC
B
GUS A3
C
GUS A4
D
GUS A5

**Fig. S6. Expanded GUS reporter assays support a role for *Brevundimonas* sp. MF374 cell-flg22 interactions in the attenuation of flg22-*Pa*-triggered plant immunity.** GUS reporter assays of MF374 wild type (WT) and deletion mutants using *A. thaliana pCYP71A12*::*GUS* seedlings to evaluate the contributions of **(A)** co-culture medium containing bacteria (A2), **(B)** washed seedlings transferred to fresh medium (A3), **(C)** cell-free co-culture supernatant (A4), and **(D)** flg22-*Pa* preincubated with bacteria followed by removal of bacterial cells (A5). Bacteria were applied at initial $OD_{600}$ values of 1 to $10^{-5}$ with 1 μM flg22-*Pa*. Fractions indicate the number of blue-stained root tips relative to the total number examined across three biological replicates. Images at $OD_{600} = 10^{-2}$ and $10^{-3}$ are also presented in **Fig. 3B**. Results for the standard co-culture assay (A1) are shown in **Fig. 2A**. NB, no bacteria; mock, sterile deionized water control.

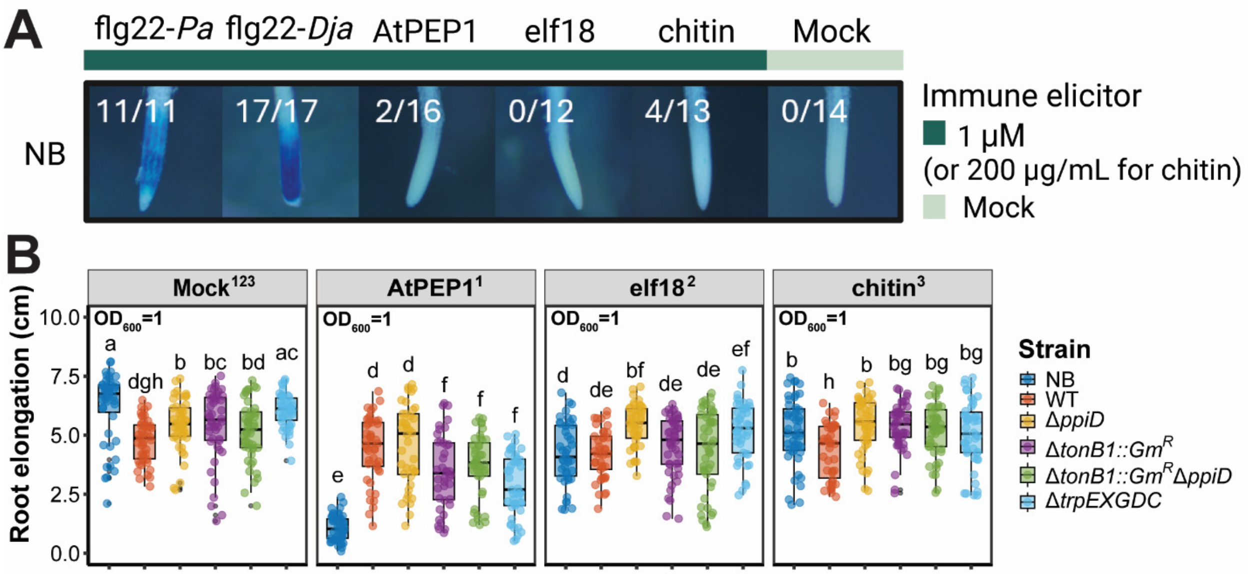


**Fig. S7. *Brevundimonas* sp. MF374 attenuates RGI triggered by AtPEP1 but not responses to elf18 and chitin under the conditions tested. (A)** GUS reporter assays of *A. thaliana pCYP71A12*::*GUS* seedlings treated with 1 µM flg22-*Pa*, flg22-*Dja*, AtPEP1, elf18 and 200 µg/mL chitin. Fractions indicate the number of blue-stained root tips relative to the total number examined across three biological replicates. **(B)** Root elongation of *A. thaliana* Col-0 seedlings inoculated with MF374 wild type (WT) or deletion mutants ($OD_{600}$ = 1), with or without 0.1 µM AtPEP1, 0.1 µM elf18, or 20 µg/mL chitin. Each treatment included three biological replicates with two technical replicates each. Different letters indicate significant differences ($P < 0.05$; Kruskal-Wallis test followed by Dunn's test with Benjamini-Hochberg correction). The same superscript number indicates multiple comparison analysis was taken within and between the elicitor group and mock group. NB, no bacteria; mock, sterile deionized water control.

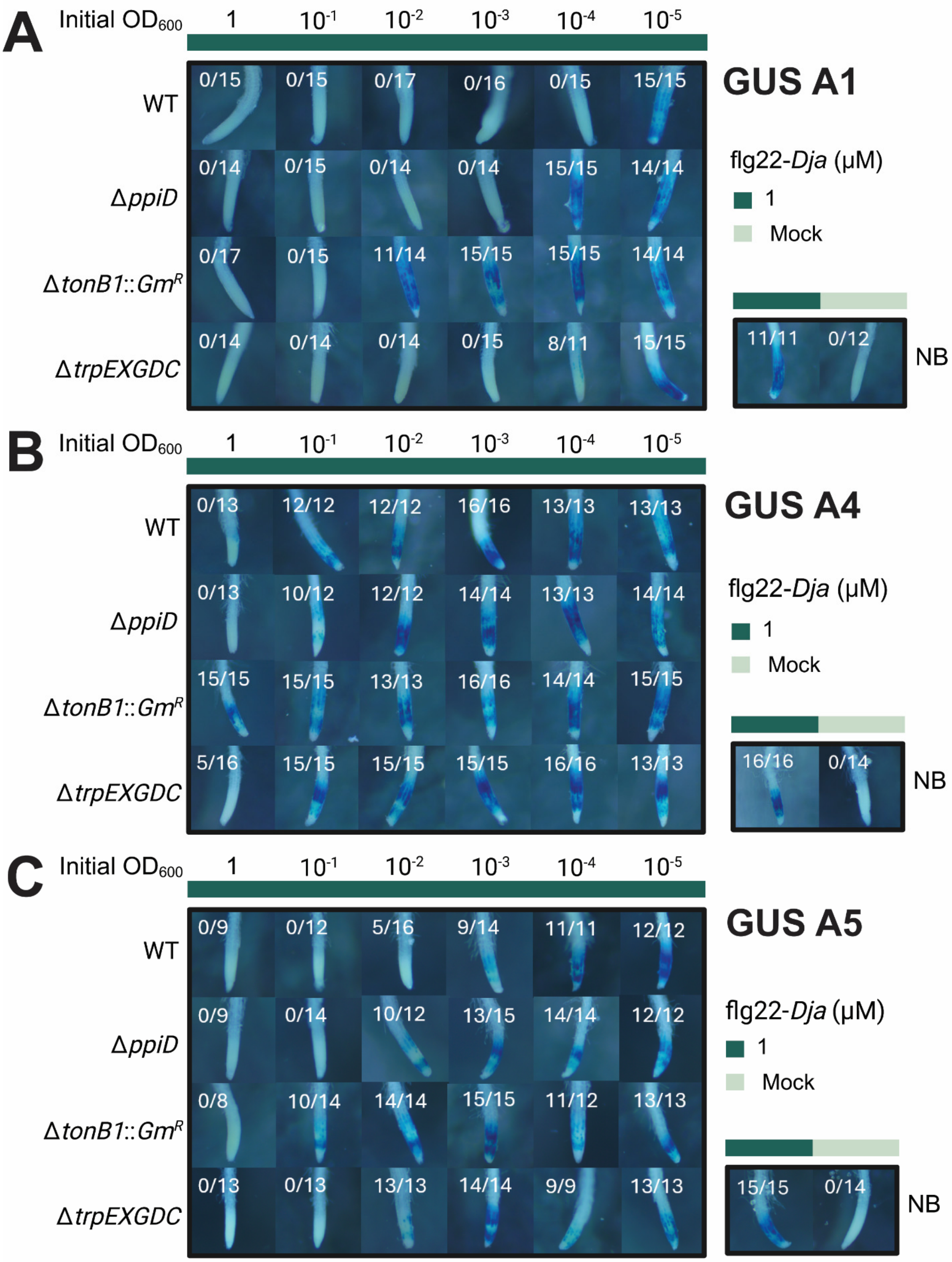
A
Initial OD600
1
10-1
10-2
10-3
10-4
10-5
GUS A1
WT
0/15
0/15
0/17
0/16
0/15
15/15
ΔppiD
0/14
0/15
0/14
0/14
15/15
14/14
ΔtonB1::GmR
0/17
0/15
11/14
15/15
15/15
14/14
ΔtrpEXGDC
0/14
0/14
0/14
0/15
8/11
15/15
flg22-Dja (μM)
1
Mock
11/11
0/12
NB
B
Initial OD600
1
10-1
10-2
10-3
10-4
10-5
GUS A4
WT
0/13
12/12
12/12
16/16
13/13
13/13
ΔppiD
0/13
10/12
12/12
14/14
13/13
14/14
ΔtonB1::GmR
15/15
15/15
13/13
16/16
14/14
15/15
ΔtrpEXGDC
5/16
15/15
15/15
15/15
16/16
13/13
flg22-Dja (μM)
1
Mock
16/16
0/14
NB
C
Initial OD600
1
10-1
10-2
10-3
10-4
10-5
GUS A5
WT
0/9
0/12
5/16
9/14
11/11
12/12
ΔppiD
0/9
0/14
10/12
13/15
14/14
12/12
ΔtonB1::GmR
0/8
10/14
14/14
15/15
11/12
13/13
ΔtrpEXGDC
0/13
0/13
13/13
14/14
9/9
13/13
flg22-Dja (μM)
1
Mock
15/15
0/14
NB

**Fig. S8. Deletion of *tonB1* impairs cell-dependent attenuation of flg22-*Dja*-triggered immunity by *Brevundimonas* sp. MF374.** GUS reporter assays of MF374 wild type (WT) and deletion mutants using *A. thaliana pCYP71A12*::*GUS* seedlings to evaluate **(A)** plant-bacteria co-culture (A1), **(B)** cell-free co-culture supernatant (A4), and **(C)** flg22-*Dja* preincubated with bacteria followed by removal of bacterial cells (A5). Bacteria were applied at initial $OD_{600}$ values of 1 to $10^{-5}$ with 1 µM flg22-*Dja*. Fractions indicate the number of blue-stained root tips relative to the total number examined across three biological replicates. Images at $OD_{600} = 10^{-2}$ and $10^{-3}$ are also presented in **Fig. 4A**. Mock-treated images for **(A, B, and C)** refer to **Fig. 2A**, **Fig. S6C**, and **Fig. S6D**, respectively. WT: MF374 wild type. NB: no bacteria. Mock: sterile deionized water control.

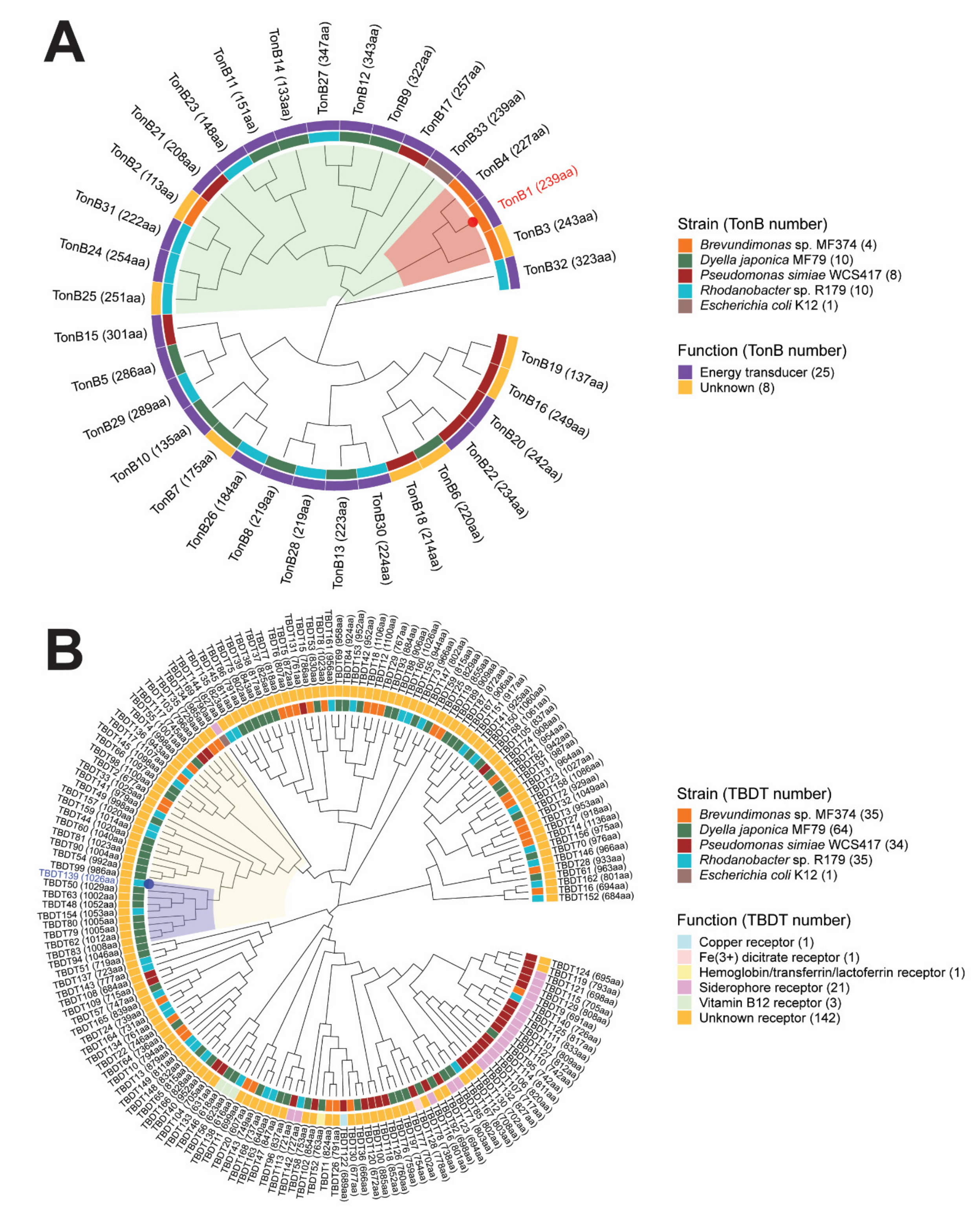


**Fig. S9. Phylogenetic analysis of TonB proteins and TonB-dependent transporters from four plant-associated immune-attenuating bacteria and *Escherichia coli* K-12. (A)** Maximum-likelihood phylogenetic tree of 33 TonB proteins from *Brevundimonas* sp. MF374, *Dyella japonica* MF79, *Pseudomonas simiae* WCS417, *Rhodanobacter* sp. Root179 (R179), and

*Escherichia coli* K-12. The red dot identifies the MF374 TonB1 (WP_017505195.1). Green and red shading indicate successively narrower clades containing MF374 TonB1. Among the TonB proteins, 25 of them were annotated as energy transducers, and the function of eight proteins was unknown based on NCBI RefSeq database. The number at the end of each TonB or TBDT was assigned based on the ascending order of the protein accession and the strain order. **(B)** Maximum-likelihood phylogenetic tree of 169 TonB-dependent transporters (TBDTs) from the same five bacterial strains. The purple dot identifies TBDT139 (WP_056714781.1), previously implicated in immune attenuation by *Rhodanobacter* sp. R179 identified by Ordon et al. (*21*). Yellow and purple shading indicate successively narrower clades containing TBDT139. In both **A** and **B**, colored annotation rings indicate bacterial species (inner ring) and predicted protein functions (outer ring). Trees were constructed using the Multiple Alignment using Fast Fourier Transform (MAFFT) algorithm and maximum-likelihood (ML) analysis method with the Jones-Taylor-Thornton (JTT) substitution model and the Nearest-Neighbor-Interchange (NNI) ML heuristic method. Protein identifiers and annotations are provided in Supplementary Table S10.

**Table S1. (separate file)**
*Brevundimonas* sp. MF374_pTGG31 and MF374_pDZ003 RB-TnSeq data summary

**Table S2. (separate file)**
Identification of the Tn-mutated genes in the five MF374 mutant candidates based on Arbitrary PCR and in the RB-TnSeq gene pool

**Table S3. (separate file)**
Bacterial strains used in this study

**Table S4. (separate file)**
Plasmids used in the RB-TnSeq library construction (*22*, *47*, *48*)

**Table S5. (separate file)**
Primers and oligonucleotides used in RB-TnSeq library construction and Tn-mutated gene identification (*19*, *22*, *44*)

**Table S6. (separate file)**
Oligonucleotides used for RB-TnSeq library construction (*42*)

**Table S7. (separate file)**
Sequence of the designed gene knockout and gene complement vectors

**Table S8. (separate file)**
Primers used for constructing the vectors of *Brevundimonas* sp. MF374 gene knockouts and gene complements

**Table S9. (separate file)**
Five immune elicitors used in this study (*5*, *17*, *21*)

**Table S10. (separate file)**
TonB and TBDT Proteins from five bacterial strains

**Table S11. (separate file)**
Characteristics of proteins in the proposed model of plant immunity attenuation by *Brevundimonas* sp. MF374
